# Pannagram: unbiased pangenome alignment and mobilome calling across insects extends horizontal transfer from transposons to polintons

**DOI:** 10.1101/2025.02.07.637071

**Authors:** Anna A. Igolkina, Aleksander Bezlepskii, Kirsten Senti, Magnus Nordborg

## Abstract

Pangenomes capture genetic variation beyond the scope of single-reference analyses, but interpreting this variation requires approaches that translate whole-genome comparisons into biologically meaningful features. We present Pannagram, a framework combining reference-free linear whole-genome alignment with feature extraction for single-nucleotide polymorphisms (SNPs), structural variants (SVs), and mobile element families (MEFs), together with a library of tools for downstream analysis. Pannagram identifies MEFs by a single criterion, recent mobility, independent of prior transposable element (TE) models or homology. Illustrating its utility through a meta-analysis of pangenome data from 60 insect species, we show that highly similar MEFs, by definition recently active, are frequently shared between species, providing strong evidence for horizontal transfer (HT) orthogonal to sequence similarity and phylogeny. We further find that vectors for such HT include not only viruses but also polintons, supported by cases of polinton HT and by polintons that remain active after insertion of TE cargo.

## Introduction

The falling cost and rising accuracy of long-read sequencing have ushered in an era of whole-genome assemblies [Logsdon et al., 2020], shifting population genetics and comparative genomics toward questions that require aligning multiple genomes. The joint representation of multiple genomes is commonly called a pangenome [Matthews et al., 2024], a vague concept spanning very different data models for tasks that range from describing variation in gene content [Tettelin et al., 2005] to fully characterizing all sequence-level polymorphisms [Liao et al., 2023].

Pangenome representations largely fall into two conceptual classes: graph-based models and linear alignment models [Kopalli et al., 2025]. In theory, graph-based pangenomes provide a flexible, reference-free framework that reduces reference bias and resolves complex genomic events. In practice, such clarity is difficult to achieve in repeat-rich regions and in regions with strong haplotype structure, where graph topologies become highly entangled [Igolkina et al., 2025]. Graph-based approaches thus perform well for genomes with low variation, such as human and many mammals [Liao et al., 2023], but their application to plant genomes remains challenging due to extensive transposable element (TE) activity and structural complexity.

In contrast, linear pangenomes offer a simpler, more naturally interpreted representation: an intuitive coordinate system, straightforward lift-over of annotations between genomes, and broad compatibility with reference-based tools. However, most current linear approaches remain anchored to a reference genome, and those that approximate reference-free representations rely on linearized projections of graph-based pangenomes [Hickey et al., 2024, Garrison et al., 2024], leaving a gap in methods for the direct construction of reference-free linear pangenomes.

We address this gap with Pannagram, a framework for direct construction of linear pangenomes without reference bias, which also extracts polymorphisms and other genomic features and is accompanied by an R package for downstream analysis (Fig. 1a). Crucially, Pannagram provides a complete workflow, from raw genomes to a reference-free coordinate system and downstream analysis of SNPs, SVs, and the mobilome, within a single integrated framework. This eliminates the need to assemble multiple independent tools and makes genome-wide, reference-free mobilome analysis practical. Its modular design keeps it flexible: the command-line interface is organized into task-specific modules, the core alignment module is a pipeline of replaceable steps, and the R library is a collection of functions operating on standard R data types rather than specialized classes, lowering the barrier for users with limited programming experience.

**Figure 1:**
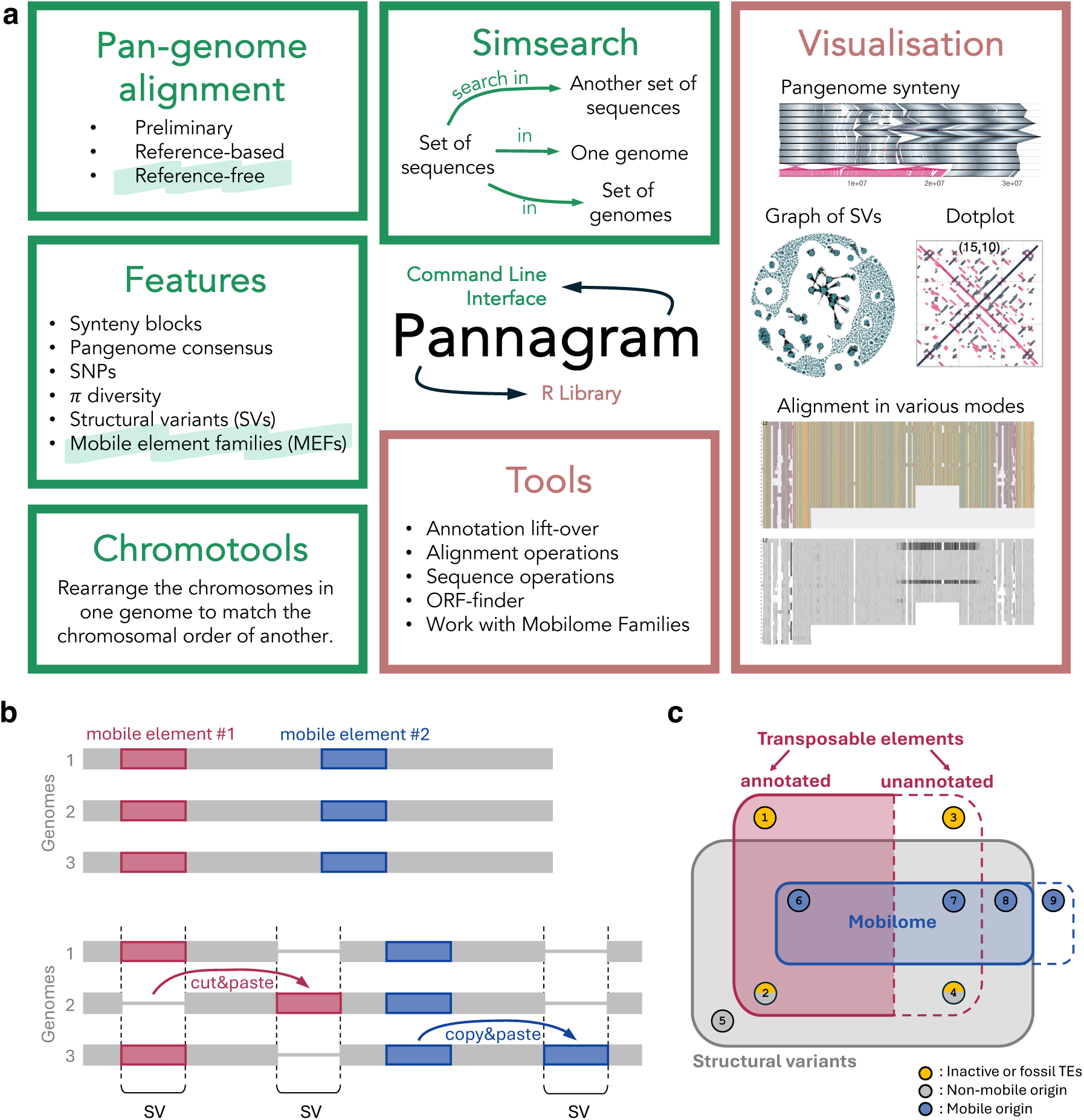
Overview of the Pannagram framework and the mobilome concept. **a**, Overview of the Pannagram framework. Pannagram comprises two main components: a command-line interface (green) and an R library (red). The command-line interface includes four functional modules (distinct tools for reference-free linear whole-genome alignment and extraction of mobile element families are highlighted). The R library provides tools for downstream analyses and visualization. A brief description of the modules and library can be found in Methods; software and full documentation is available on GitHub. **b**, Mobile elements generating SVs in the whole-genome alignment via “cut & paste” and “copy & paste” mechanisms. **c**, The relationship between the mobilome, TEs, and SVs. The mobilome comprises genomic elements with evidence of mobility, as reflected by their presence in SVs. It includes both known and previously unannotated TEs, as well as mobile elements of non-TE origin. Unannotated TEs are either non-autonomous, highly diverged from known TEs, or belong to previously uncharacterized families. (1) Annotated TEs that have lost mobility and are not represented in SVs. (2) Known TEs detected in SVs may be either active or inactive, representing TE fragments. (3–4) Same as A and B, respectively, but for previously unannotated TEs. (5) SVs of non-TE and non-mobile origin. (6) Known TEs within the mobilome. (7) Previously unannotated TEs within the mobilome. (8) Mobilome components of non-TE origin. (9) Mobile sequences lacking evidence in SVs therefore excluded from the mobilome under our definition.

Reference-free pangenomes provide a near-complete representation of genomic variability, in particular structural variants (SVs) that are poorly captured by mapping short reads to reference genomes [Mahmoud et al., 2019]. Definitions of an SV differ; here we consider a locus with alternative length variants observed at homologous positions in the pangenome coordinate system, from 1-bp indels onward (inversions and intrachromosomal translocations are also identifiable from Pannagram alignments but not directly called by the Features module). Pannagram provides tools for focused analysis of the most dynamic category of SVs: the mobilome (Fig. 1b).

We define the mobilome as the set of genomic sequences with evidence of mobility, using a single criterion: a sequence must recur within the full set of SVs as a group of related variants in different places in the genome, forming what we call a “mobile element family” (MEF) (Fig. 1c). Because our approach does not rely on prior assumptions about TE structure or sequence homology, MEFs are defined solely by shared mobility and need not correspond to conventional TE families. Consequently, they include elements typically missed by standard TE annotation, such as non-autonomous elements lacking detectable homology to autonomous counterparts, as well as potentially novel TE classes (Fig. 1c). Conversely, “fossilized” TEs that no longer exhibit presence–absence polymorphism are excluded. The approach relies on sufficient sample size: when few genomes are available, low-activity families may be observed too rarely to be inferred as MEFs. A related method, pantera, similarly infers mobile-element families from pangenome polymorphism, though from graph rather than linear reference-free alignments [Sierra and Durbin, 2024].

The utility of Pannagram for mobilome analysis was recently illustrated in a pangenome study of *Arabidopsis thaliana* [Igolkina et al., 2025]. Here we describe Pannagram in detail while performing a comparative analysis of insect pangenomes and mobilomes. Unlike classical comparative genomics, which directly compares genomes between species, we first characterize within-species variation and then compare this variation across species, extending pangenomes from their usual within-species use to cross-species comparison of polymorphisms. We focus in particular on horizontal transfer (HT), which is well documented in insects [Peccoud et al., 2017, Reiss et al., 2019, Li et al., 2022, Scarpa et al., 2024]. Pannagram allows us to use the co-occurrence of near-identical, active MEFs in different species as evidence for HT that is independent of the sequence-similarity and phylogenetic arguments on which HT inference typically relies. Beyond identifying candidate event, we describe potential molecular mechanisms for HT. Two of our findings appear to be previously unreported: recent HT of polintons between distantly related insects and mobile polintons carrying transposable-element cargo. Both extend earlier reports of deeper-time polinton host-switching [Widen et al., 2023, Jeong et al., 2023] and of Lepidoptera as an HT hub [Reiss et al., 2019]. Our analysis demonstrates the capabilities of Pannagram and reveals how pangenomes themselves can shape questions on mobilome dynamics at an interspecific level.

## Results

### Structure of the Pannagram package

Pannagram consists of two components installed together (Fig. 1a): a command-line interface (CLI) for linear whole-genome alignment, feature extraction, and similarity search, and an R library for downstream analysis and visualization. Full documentation is available at GitHub.

The CLI includes four modules (see Methods). The Pannagram module performs whole-genome alignment in three modes—preliminary, reference-based, and reference-free—each producing a matrix of corresponding chromosomal positions (negative values encode inversions). The Features module, which requires a Pannagram alignment, extracts synteny blocks, SNPs (VCF), SVs (GFF and FASTA), and MEFs derived from SVs (Fig. 2a). The Chromotools module cuts and rearranges query chromosomes to match a reference, fixing large-scale assembly errors and arbitrary differences in chromosome numbering and orientation. The independently run Simsearch module searches a set of sequences (e.g., SVs) for matches in another set of sequences or genomes, by identity and coverage.

**Figure 2:**
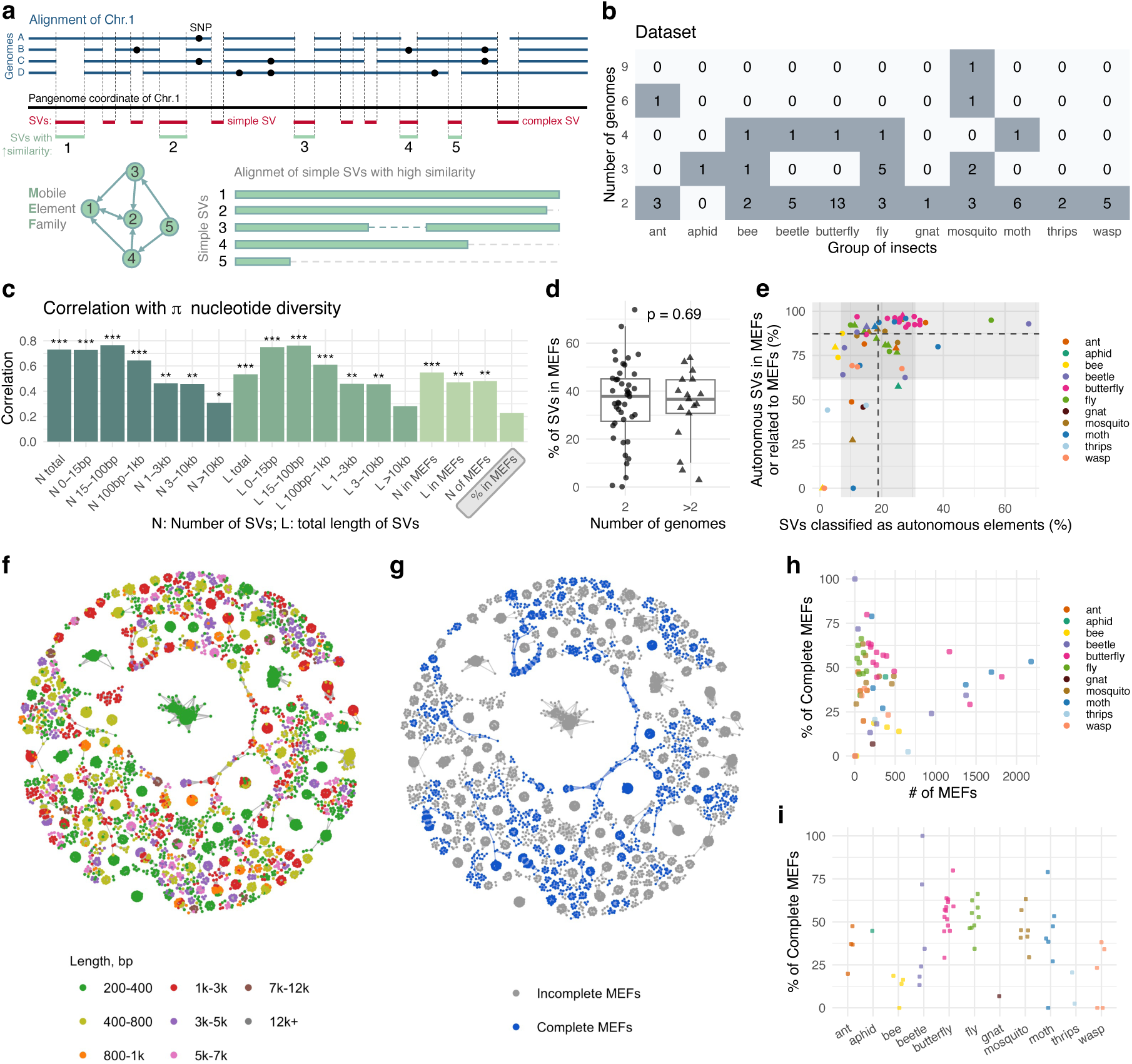
Meta-analysis of whole-genome polymorphisms. **a**, Pipeline for extracting mobile element families (MEFs) from whole-genome alignments. SVs are filtered to retain simple events, which are then compared and clustered into graphs of nestedness. **b**, Dataset composition, showing the distribution of species across insect groups according to the number of available genome assemblies. **c**, The correlation between SV-derived metrics and nucleotide diversity (*π*), restricted to species represented by two genome assemblies, is weaker for longer elements, suggesting distinct population dynamics. N denotes the number of SVs, and L their total length. The grey box highlights that the percentage of SVs in MEFs does not correlate with *π*. **d**, The proportion of SVs assigned to MEFs does not appear to depend on the number of genomes analysed. **e**, Fraction of SVs classified as autonomous elements and fraction of these associated with MEFs. Dashed lines, mean; grey shaded areas, ± s.d. **f–g**, Graph-based representation of MEFs in *Anopheles arabiensis* (mosquito), illustrating (f) size distribution and (g) completeness, i.e., the presence of autonomous elements. The same graph, with nodes colored according to the presence of autonomous elements, is shown in Extended Data Fig. 1a. **h–i**, Variation in the number and complete MEFs across insect groups.

The R library supports post-analysis of SVs and MEFs, including annotation lift-over, dot plots, ORF prediction and visualization, multiple sequence alignment display, and MEF graphs. All visualizations presented in this paper were generated using this library.

### Benchmarking

We benchmarked Pannagram against established tools of the same class on cost, coverage and accuracy; full per-point measurements, commands and simulation protocols are given in the Supplementary Information.

#### Reference-free multiple-genome alignment

We compared Pannagram’s reference-free alignment against three tools of the same class: Progressive Cactus v3.2.1 [Armstrong et al., 2020], pro-gressiveMauve build 2015-02-13 [Darling et al., 2010], and SibeliaZ v1.2.7 [Minkin and Medvedev, 2020]. Because SibeliaZ’s base-level mode could not run at these scales, exceeding 256,GB of memory, it was evaluated only as a synteny-block method. We used intraspecific chromosome-scale panels of *Drosophila melanogaster* (∼135,Mb, 2–11 genomes) and *Anopheles coluzzii* (∼247,Mb, 2–8 genomes), with all tools receiving byte-identical inputs and running on 16 cores. Peak memory was the clearest differentiator: Pannagram used 3.6–16.0,GB, an order of magnitude below Cactus (24.5–46.8,GB) and, at scale, below Mauve (up to 31.3,GB), while still emitting a base-level alignment with per-site SNP and SV calls. On 11 *D. melanogaster* genomes it finished in 51.3,min against 2.5,h for Cactus and 10.8,h for Mauve, scaled approximately linearly with the number of genomes, and aligned a fraction of each genome comparable to Cactus (0.94 vs 0.93). On simulated genomes with known homology (simuG; *E. coli*, *D. melanogaster*, and *Anopheles* chromosomes), Pannagram matched Cactus on SNPs (*F*_1_ ≥ 0.997 throughout) and was the best or tied-best method on inversions at every point, uniquely showing no degradation with divergence (*F*_1_ = 0.998 to 1.000); progressiveMauve lost 20–40% of homologous bases on inversions.

#### Mobile-element families vs de-novo TE annotation

To test what the pangenome view of the mobilome recovers beyond de-novo TE annotation, we compared Pannagram’s MEFs with EDTA (v2.3) [Ou et al., 2019] on the five-genome *Drosophila melanogaster* panel. Pannagram grouped 771 large SVs into 57 MEFs, of which 19 (321 of 771 members) were absent from EDTA’s *de novo* annotation by both sequence and coordinate. 17 of the 19 map to named *D. melanogaster* elements, strongly enriched for non-LTR LINEs (Jockey, I, R1, Loa) and Tc1-Mariner DNA transposons, the classes that *de novo* structural annotators resolve worst; the remaining two are short (≤300 bp) and unclassified. These 17 MEFs are genuine, canonical, polymorphic TEs: 263 of 270 members are recovered as their named element by RepeatMasker [Smit et al., 2013–2015] with the curated library, so their absence reflects an incomplete *de novo* library rather than undetectability. Another reason Pannagram may recover more elements is that it counts copies along the pangenome coordinate rather than within genomes independently. An element present once per genome but at different loci appears as a singleton to *de novo* annotation, yet as multiple SVs across genomes, enough to define an MEF.

#### Simsearch vs RepeatMasker

Pannagram’s Simsearch addresses the same task as Repeat-Masker, locating library sequences within genomes and reporting copy number. Against simulated ground truth (21 genomes, 0–30% divergence; an 816-sequence *A. thaliana* library implanted in a homology-free background), Simsearch matched RepeatMasker on the near-full-length copies it targets up to 15% divergence (recall 0.993 vs 0.999; paired Δ = −0.005, bootstrap 95% CI [−0.012, +0.001], *p* = 0.15) at precision ≥ 0.9999, at 5–7× lower wall-clock and an order of magnitude less CPU-time, with comparable memory (∼0.2 GB). On real cross-species sequence (the same library searched against *A. lyrata*, 207 Mb), every locus and base pair Simsearch reports is also reported by RepeatMasker (100% containment), making it precision-perfect against the community standard without a ground truth, and 9× faster; its default 85–85 thresholds (85% identity and coverage) report recent, near-full-length copies and are tunable to recover the divergent, fragmentary tail (for details, see Supplementary Information).

### Meta-analysis of whole-genome polymorphism in insects

The standard Pannagram pipeline for extracting whole-genome polymorphisms is as follows (Fig. 2a). The Pannagram module first constructs a per-chromosome linear reference-free alignment that resolves inversions and translocations, defining a “pangenome” coordinate system that connects all genomes and facilitates annotation lift-over. The Features module then derives a “consensus sequence” (containing *all* presence alleles) with SNPs and SVs in pangenome coordinates, projected onto individual genomes. The consensus nucleotide at each position is the most frequent cunleotide. SVs are subdivided into simple (biallelic presence/absence) and complex (three or more alleles). SV sequences are extracted from the consensus, and the module uses Simsearch to compare simple SVs and build MEF graphs from similarity and coverage (e.g., a standard 80–80 rule for TEs [Wicker et al., 2007]), whose members can be aligned to characterize nestedness (Fig. 2a).

We collected all available insect whole-genome assemblies from GenBank, restricting ourselves to genomes not exceeding 300 Mbp to minimize genome-size biases. In total, 60 species were included (Supplementary Table 1). Of these, 43 were represented by only two independent assemblies (Fig. 2b), in most cases the two haplotypes of a single diploid individual, reflecting still-limited sampling. For outcrossing species under random mating, divergence between these haplotypes provides an unbiased estimate of nucleotide diversity (*π*), allowing cross-species comparison of SVs and mobilome relative to background diversity.

Across species, the number of short SVs is strongly correlated with *π*, suggesting both reflect neutral or nearly neutral evolution and an underlying “effective population size”. This correlation weakened with increasing SV length, however, indicating that larger-scale events follow different dynamics, as expected for mobile element insertions (Fig. 2c). The fraction of SVs forming MEFs (each a connected component of the MEF graph) does not correlate with *π*, phylogenetic relatedness, or the number of genomes analysed (Fig. 2c–d), likely reflecting species-specific mobile element activity decoupled from overall genomic diversity.

Distinguishing autonomous from non-autonomous elements generally requires functional experiments; as a proxy, we classified as putatively autonomous only SVs encoding ORFs similar to known Repbase ORFs (see Methods) [Bao et al., 2015], which likely overestimates autonomy since not all ORFs are functional. Nonetheless, 10–30% of SVs were classified as autonomous, and nearly all (87%) were part of or related to MEFs, supporting MEFs as biologically meaningful units (Fig. 2e). Although ORFs in autonomous MEFs were of Repbase-like length, they showed limited similarity: among 12,514 autonomous MEFs, average similarity to the closest Repbase match was 30% and only 27 exceeded 90% (Extended Data Fig. 2a), indicating that Repbase captures only a small fraction of the global mobilome.

Examples of MEF graphs for mosquito *Anopheles arabiensis* are shown in Fig. 2f–g. These graphs reveal substantial heterogeneity in MEFs: many MEFs consist only of short, likely non-autonomous sequences, whereas others contain longer, likely autonomous elements. We accordingly classified MEFs as “complete” or “incomplete” by the presence of an autonomous element. Incomplete MEFs may arise from high non-autonomous mobility with low activity of the corresponding autonomous element, or from reliance on autonomous elements from other MEFs without high similarity, analogous to Alu mobilization by LINE retrotransposons in humans [Ahl et al., 2015]. The median proportion of complete MEFs is 45% (Fig. 2h) but varies considerably between species (Fig. 2i). To test whether this variation could stem from Repbase annotation bias, we screened long ORFs (≥ 200 aa) from incomplete MEFs against SwissProt [UniProt Consortium, 2025], Pfam [Mistry et al., 2021], and UniRef90 [Suzek et al., 2015] (excluding spurious low-complexity ORFs). This identified 32 additional TE-related MEFs but left the among-group patterns unchanged, arguing against an annotation artefact (Extended Data Fig. 1). We also found 74 MEFs lacking TE genes and carrying only protein-coding cargo, indicating ongoing gene duplication mediated by transposon machinery (Extended Data Fig. 1e).

### Horizontal transfer of mobile elements

MEFs from different species were compared at 95% identity and 90% coverage, more restrictive than the thresholds defining MEFs (85–85), to identify highly similar MEFs whose cross-species conservation is difficult to explain by long-term vertical inheritance. As markers of recent mobility, such shared MEFs provide evidence for HT beyond sequence similarity alone. The number of shared MEFs was markedly higher between sister species (Fig. 3a), consistent with vertical inheritance or recent hybridization, but conserved MEFs were also detected between phylogenetically distant species, including moths and butterflies, which diverged nearly 100 Myr ago [Kawahara et al., 2019]. Putative transfers included both Class I and Class II TEs (Extended Data Fig. 2b). Only three of 1,217 shared MEFs were previously in Repbase, all involving closely related congeners (*BEL7-I AG* and *CR1-5 AG*, annotated in *A. gambiae* but active in *A. arabiensis* and *A. coluzzii* ; and *Gypsy-15 HeAr*, active in both *H. armigera* and *H. zea*), making HT less distinguishable from vertical inheritance than in the distant-species examples below.

**Figure 3:**
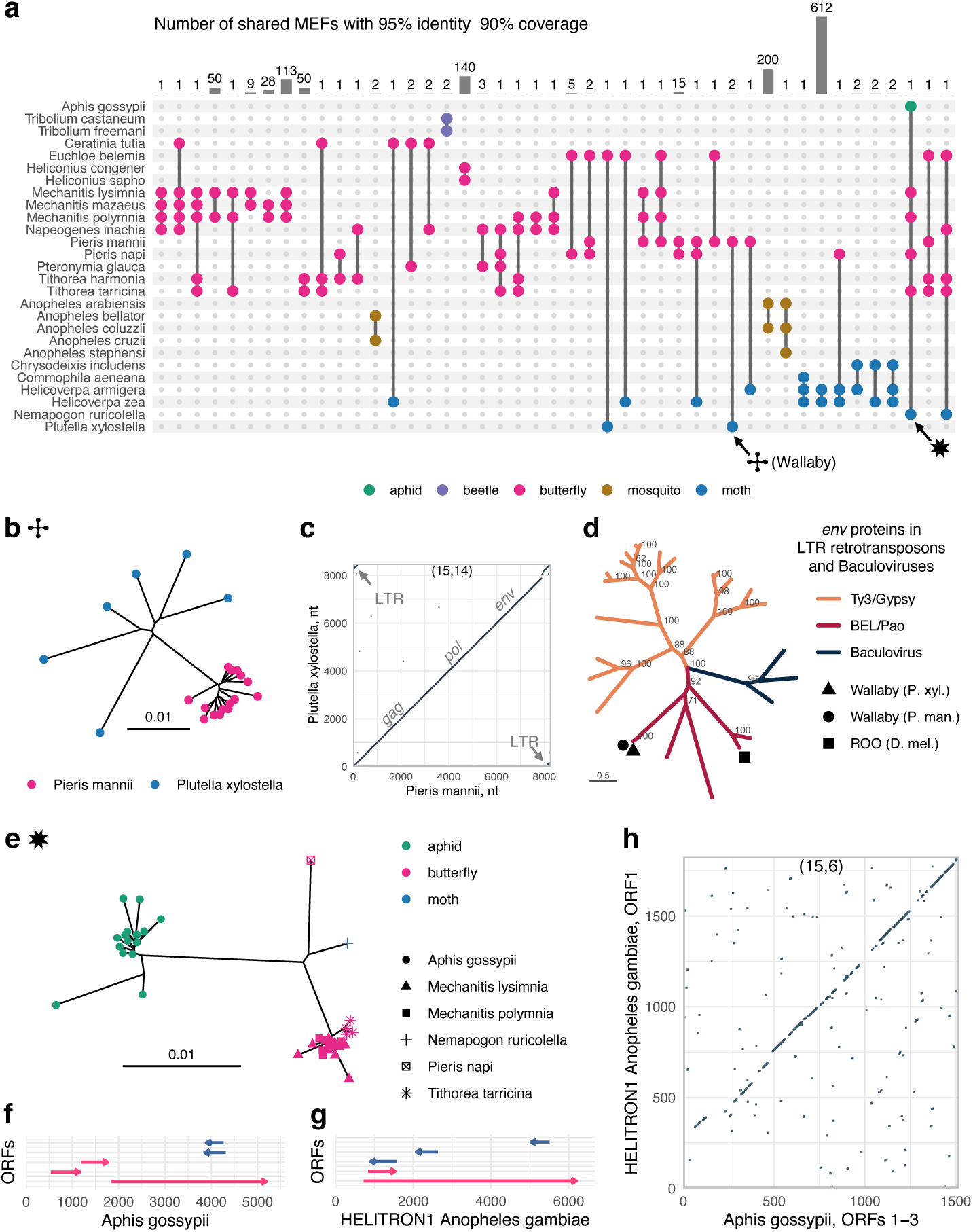
Horizontal transfer of TE. **a**, UpSet plot showing the number of shared mobile element families (MEFs) across insect species at 95% sequence identity and 90% coverage. Connected dots indicate the species sharing MEFs; bars indicate the number of shared MEFs for each species combination; colours denote insect groups. **b–c**, Example of HT of the MEF marked by a cross in **a**. **b**, Neighbour-joining tree of nucleotide sequences of the shared MEFs. **c**, Dot plot of representative nucleotide sequences from MEFs in *Pieris mannii* (butterfly) and *Plutella xylostella* (moth). Parameters for dotplot: window size 15; number of matches 14. **d**, Phylogenetic tree of the *env* gene from the representative *Wallaby* MEF sequence and well-characterized LTR retrotransposons, placing *Wallaby* in a distinct branch of the Belpaoviridae (Bel/Pao) family, distant from the *roo* element of *D. melanogaster*. **e–h**, Example of HT of the MEF marked by a star in **a**. **e**, Neighbour-joining tree of nucleotide sequences of the shared MEFs. **f**, ORF organization of the representative MEF sequence in *Aphis gossypii* (aphid). **g**, ORF organization of the helitron from *Anopheles gambiae* (*HELITRON1 AG* in Repbase). **h**, Dot plot of the concatenated amino acid sequence from three ORFs of the representative *A. gossypii* element and the longest ORF from the helitron of *Anopheles gambiae*. The *A. gossypii* sequences do not fully align with the helitron, suggesting that this element might be non-autonomous.

One such shared MEF, classified as a long terminal repeat (LTR) retrotransposon, was found to be shared between the moth *Plutella xylostella* and the butterfly *Pieris mannii* (Figs. 3b–d); the mean divergence between species is 0.32 based on BUSCO genes [Seppey et al., 2019] (Extended Data Fig. 3). This element, which we name *Wallaby*, encodes a single 2,475-aa ORF (Extended Data Fig. 4a), its closest Repbase match (33% aa identity) being the 2,377-aa *gag* -*pol* -*env* polyprotein of the *roo* element in *Drosophila melanogaster* (*ROO I* ; Extended Data Fig. 4b,c) [de la Chaux and Wagner, 2009]. Phylogenetic analysis of its *pol* and *env* regions with well-characterized LTR retrotransposons [Malik et al., 2000, Malik and Henikoff, 2005, Senti et al., 2026] placed *Wallaby* in a distinct branch of the Belpaoviridae (Bel/Pao) family [Soriano et al., 2021], but distant from *roo* (Fig. 3d; Extended Data Fig. 4d). The *Wallaby env* shares characteristic features with envelope proteins of DNA baculoviruses and infectious endogenous retroviruses of the Ty3/gypsy clade [Senti et al., 2026, Pearson and Rohrmann, 2006]—an N-terminal signal peptide, furin cleavage site, fusion peptide, conserved cysteines, and a transmembrane domain [Westenberg et al., 2002, 2004] (Extended Data Fig. 5). Although direct evidence for cell-to-cell infectivity of these Bel/Pao retroelements is lacking, this suggests that *Wallaby*, when expressed, may form extracellular viral particles bearing envelope glycoproteins capable of entering recipient cells, a property proposed to facilitate horizontal transfer between hosts.

A second example is a MEF shared among six distant species, including butterflies, moths, and an aphid (Fig. 3a,e). Its ORFs resemble the Repbase helitron *HELITRON1 AG* from the mosquito *Anopheles gambiae* (Figs. 3f–h), which encodes a replication–helicase fusion protein [Kapitonov and Jurka, 2003] but, unlike *Wallaby*, lacks an envelope protein.

These results add to the growing evidence that horizontal transfer of transposons is common in insects [Reiss et al., 2019, Bargues and Lerat, 2017]. The mechanisms remain poorly understood, especially for elements lacking an obvious means of cell-to-cell transmission and thus requiring external mediators—which the rest of this paper examines.

### Horizontal transfer of polintons

Of the 2,520,364 SVs identified across insect species, 69,134 exceeded 1 kb (the usual lower size threshold for TEs) and 8,360 exceeded 12 kb (above the typical TE size). Clustering these SVs by the similarity of their ORFs (≥150 aa) partitioned 4,224 SVs into two clusters: “Many ORFs”, with substantially more ORFs per SV, and “Few ORFs” (Fig. 4a; Extended Data Fig. 6a). The “Many ORFs” cluster comprised 685 SVs, with the largest contribution from moths and butterflies (Fig. 4b).

**Figure 4:**
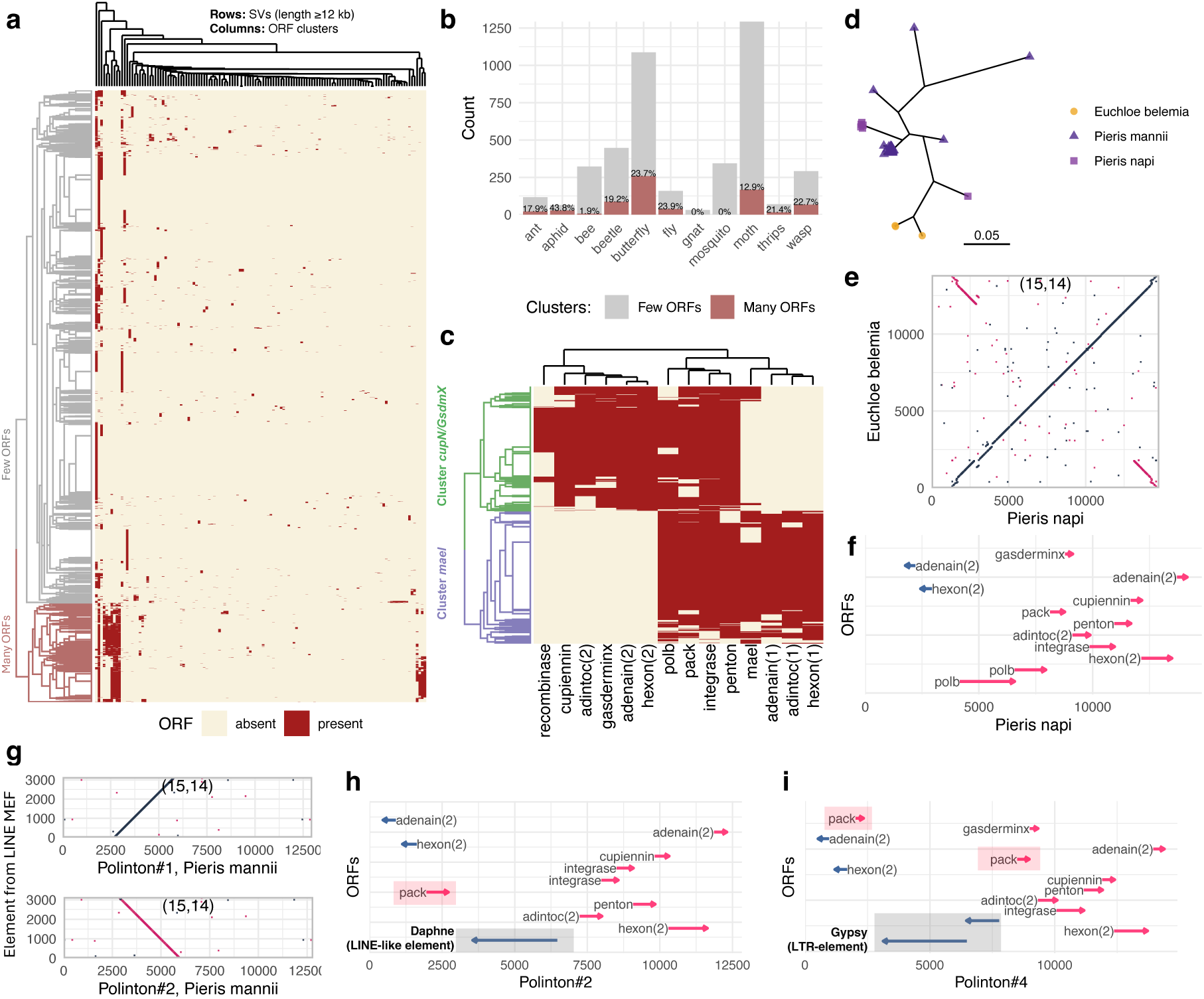
Horizontal transfer of polintons and the gene organization of polinton-related SVs. **a**, Hierarchical clustering of ORFs in SVs of length ≥12 kb into two major groups, denoted as “Many ORFs” and “Few ORFs”. **b**, Distribution of SVs from the two clusters across insect taxa. Lepidoptera exhibit a higher abundance of long SVs compared to other taxa, with a substantial fraction belonging to the “Many ORFs” cluster. **c**, The “Many ORFs” cluster comprises polintons forming two major groups: *mael* -associated and *cupiennin/gasderminX* -associated. **d**, Neighbour-joining tree of closely related polintons from *Euchloe belemia* (3 SVs), *Pieris mannii* (59 SVs), and *Pieris napi* (6 SVs). **e**, Dot-plot comparison between polintons of *P. napi* and *E. belemia* (window size 15, minimum matches 14). Terminal inverted repeats (TIRs) indicate that the corresponding SVs represent insertion events. **f**, Gene organization of the polinton from *P. napi*. **g**, Dot-plot comparisons of two polinton-related SVs (Polinton#1 and Polinton#2) from *Pieris mannii* with the SV of a LINE-associated MEF (window size 15; minimum matches 14). **h**, gene organization of the *P. mannii* polinton-related SVs carrying LINE (Daphne) cargo, which lack *polB*. **i**, gene organization of polinton-related SVs in which *polB* is replaced by a *gypsy* LTR element.

A BLAST analysis of ORFs from the “Many ORFs” cluster showed that many SVs encode hallmark polinton genes—polymerase (*polB*), *integrase*, *penton*, *pack* (packaging ATPase), *hexon*, and *Adintoc*—indicating a polinton origin [Kapitonov and Jurka, 2003, Pritham et al., 2007], and also encode *adenain*, consistent with classification as adintoviruses (Fig. 4c) [Koonin et al., 2024, Starrett et al., 2021].

Within this cluster, two subclusters could be distinguished by allelic variants of *hexon* and *Adintoc* and by linkage either to *mael* or to the *cupiennin* and *gasderminX* (*cupN/GsdmX*) genes (Fig. 4c), with butterflies enriched for *cupN/GsdmX* and moths for *mael* (Extended Data Fig. 6b). However, clustering of the major capsid protein *hexon* shows that individual species can carry polintons from both subclusters, consistent with horizontal transfer between butterflies and moths (Extended Data Fig. 6c).

Using a conservative 90% similarity and coverage threshold, we identified one set of polinton-related SVs shared by three butterfly species: two sister species and one more distantly related species (Fig. 4d). Mean sequence divergence between *Euchloe belemia* and *Pieris napi* was 0.21 (Extended Data Fig. 3). Dot plots comparing SVs from *E. belemia* and *P. napi* revealed ∼2-kb terminal inverted repeats (TIRs), confirming genuine insertions rather than isolated polinton-like sequences (Fig. 4e). Unlike the short, non-coding TIRs of most DNA transposons, these TIRs contain genes encoding the core polinton proteins *hexon* and *adenain*, resulting in two copies of each gene per insertion (Fig. 4f).

The presence of highly similar, full-length polinton insertions in distantly related butterflies strongly supports recent horizontal transfer.

### Transposons as polinton cargo

Large polinton/maverick elements are known to transport other sequences [Widen et al., 2023]. Consistent with this, among the polinton-related SVs we identified cases that not only encode canonical polinton proteins but also contain sequences highly similar to other MEFs, likely representing “cargo”. For example, in *Pieris mannii* we found two polinton-related SVs containing identical 3.2-kb cargo sequences that differed only by internal rearrangements, consistent with a shared origin (Fig. 4h; Extended Data Fig. 7b). The cargo showed 95% identity to a highly active MEF supported by 669 SVs (Fig. 4g), classified as a LINE element: alignment of its SVs revealed a 3*^′^*-end anchoring pattern characteristic of target-primed reverse transcription [Luan et al., 1993] (Extended Data Fig. 7a), and its ORF showed homology to the Daphne LINE clade in RepBase (27% aa identity over ∼855 aa to *Daphne-5 CydSpl* from the moth *Cydia splendana*). Thus an active MEF inserted into a polinton, which then amplified.

Curiously, these two polintons also lack *polB* (Fig. 4h), a gene typically required for autonomous polintons. In contrast, *polB* is retained in other highly similar, more complete *P. mannii* polintons (Extended Data Fig. 7c–e), raising the question of how the *polB* -lacking elements are mobilized. The LINE insertion coincides with the loss of *polB* ; however, because it lies on the opposite side of the *pack* gene, the precise sequence of these events remains unclear.

A very similar arrangement was found in another group of polinton-related SVs, in which *polB* was replaced by an LTR element (closest RepBase match: *Gypsy-1 CerCap-LTR* from the fly *Ceratitis capitata*, with 49% aa identity across 1,191 aa; Fig. 4i). The high similarity within this group suggests that these elements also remained active after the LTR insertion and the loss of *polB* (Extended Data Fig. 7f–h).

All polinton-related SVs shown here are flanked by TIRs, consistent with insertion-derived elements. Overall, these examples support the idea that polintons can carry TEs while retaining activity and can also transfer them, possibly between species.

### Horizontal transfer of TEs via virus vectors

A final potential mediator of TE horizontal transfer is viruses [Gilbert et al., 2016, 2014, 2010, Fraser et al., 1985, Muller et al., 2021, Drezen et al., 2017]. Using Simsearch, we aligned all MEF SVs against the 542 Nucleopolyhedrovirus (NPV; Baculoviridae) genomes longer than 50 kb in GenBank. One MEF from the moth *Helicoverpa zea* and one from the butterfly *Euchloe belemia* (mean divergence 0.28; Extended Data Fig. 3) matched the same 436-nt region in all available *H. zea* nucleopolyhedrovirus (HzNPV) genomes (AF334030.1, KJ004000.1, KM596835.1; positions 36,011–36,446 in KM596835.1; Fig. 5a). This region is almost identical to an *H. zea* SV (3 SNPs; pointer 1) and shares the same short indel (pointer 2), consistent with a recent common origin; *E. belemia* SVs are more distant but some cover the entire region (pointer 3). The map of shared SNPs indicates a common origin of the two MEFs and the HzNPV region.

**Figure 5:**
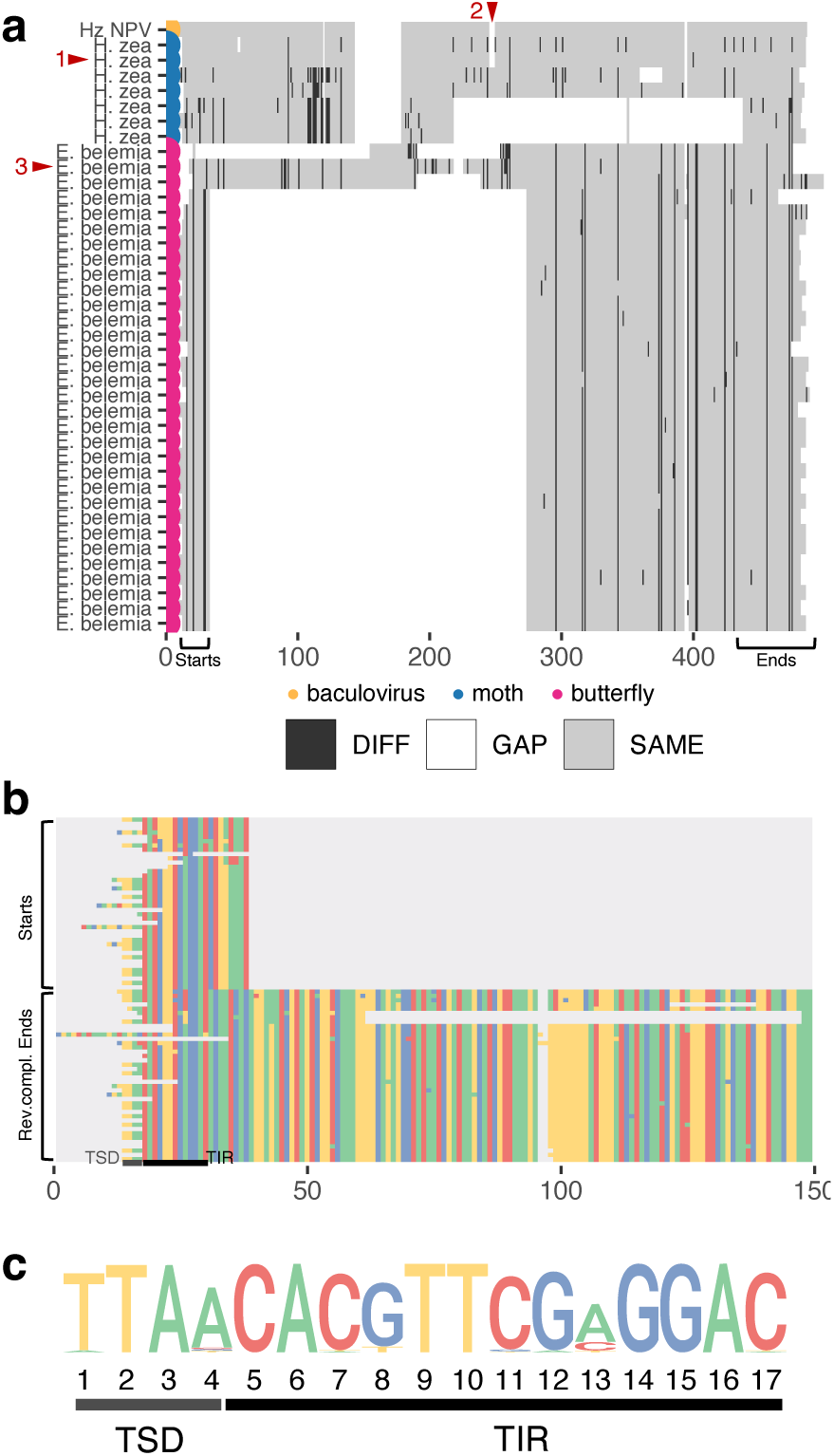
Virus-mediated transfer of TEs. **a**, Multiple sequence alignment of regions from the *Helicoverpa zea nucleopolyhedrovirus* (HzNPV) genome (yellow), sequences of a MEF from *Helicoverpa zea* (blue), and sequences of a MEF from *Euchloe belemia* (pink). Grey indicates nucleotides identical to those in the top row, black indicates differences, and white indicates gaps in the alignment. Sequence (1) from *H. zea* is almost identical to a 436 bp region in HzNPV; both also share the indel (2). The MEF from *E. belemia* contains variants of different lengths, including sequence (3), which spans the entire corresponding region in HzNPV. The start and end regions of the alignment are marked and conserved across nearly all sequences. **b,** Multiple sequence alignment of the starts and the reverse complement ends. Colours represent nucleotides. **c,** Motifs of the TSD and 13 nt TIR.

Both MEFs consist of different length variants, but the alignment shows nearly perfect matches at their starts and ends (Fig. 5a). Aligning the starts and reverse-complemented ends revealed a “TTAA” target site duplication (TSD), present only at one end of each SV, and a 13-bp terminal inverted repeat (TIR) motif at both ends (Figs. 5b–c). Such TSDs and TIRs are characteristic of the *piggyBac* DNA-transposon superfamily, indicating that these MEFs comprise non-autonomous elements, which we name *piggyMITE* (a *piggyBac*-like miniature inverted-repeat transposable element, MITE), mobilized by this superfamily [Mitra et al., 2008]. Scanning the HzNPV, *H. zea*, and *E. belemia* genomes for *piggyBac* transposases linked to the TIR motif revealed candidate autonomous elements in both hosts, though their TIRs differed from the inferred motif by several nucleotides (see Methods and Extended Data Fig. 8). These could in principle mobilize the MEFs, but the precise autonomous element remains unclear and may have escaped detection through low population frequency or loss.

The high similarity between the *H. zea* MEF and the HzNPV region is parsimoniously explained by activity of the same non-autonomous element in both host and virus, consistent with the known movement of mobile elements between host and virus during infection [Gilbert et al., 2014, Muller et al., 2021, Loiseau et al., 2021, Lerch and Friesen, 1992]. That the MEF apparently transferred between *H. zea* and *E. belemia* also resembles HzNPV suggests this virus, or a related baculovirus, acted as an HT vector.

## Discussion

Although a pangenome is conceptually the complete repertoire of variation within a population, in practice it is constrained by the available data. At the same time, even two genomes (*e.g.*, from a single outcrossing individual) can expose patterns inaccessible in a single-reference framework. Pannagram combines linear whole-genome alignment with feature extraction and downstream analysis. A key advantage is its reference-free identification of mobile element families (MEFs), supported by lightweight tools for their analysis and visualization. Benchmarking showed that Pannagram recovers real, polymorphic elements that other de-novo annotation missed. When applied across diverse insect species, it enabled analyses ranging from global patterns of genomic variation to detection of HT of transposons and polintons.

Our meta-analysis of insects showed that nucleotide diversity (*π*) is strongly associated with short SV diversity but that this relationship weakens with increasing SV length (Fig. 2c), suggesting that SNPs and short SVs are governed by related mechanisms whereas long SVs are not. Consistent with this, about 40% of long SVs are linked to mobile element activity, which may follow episodic, lineage-specific dynamics. Thus SNP-based diversity captures only part of genomic variability, and resolving pangenomic diversity requires integrating all layers of variation in a unified framework.

Horizontal transfer of transposable elements is documented between many hosts, yet how it proceeds mechanistically, from the germline of one host to that of another, remains debated and has rarely been reproduced experimentally. Most MEFs and TEs (most Class I retroelements and virtually all Class II DNA elements) replicate intracellularly in the host germline, so their transfer between the germlines of two highly diverged hosts seems unlikely without additional capabilities or biological agents.

Pannagram provides an orthogonal line of evidence for HT by identifying shared recent activity directly from presence–absence polymorphism, complementing the traditional signatures of sequence similarity and phylogenetic incongruence. Shared active MEFs across both closely and distantly related taxa, including moths, butterflies, and aphids, are therefore consistent with recent mobility in multiple lineages. The potential recency of such transfers is underscored by transposons documented to have spread horizontally between *Drosophila* species within the last few decades [Scarpa et al., 2025]. Even with limited sampling and conservative criteria for shared MEFs, this approach recovered three testable routes of HT: (i) autonomous transfer by *env* -encoding elements, (ii) transfer mediated by cargo-carrying polintons, and (iii) transfer mediated by baculoviruses (Fig. 6). These three scenarios are all plausible and have empirical support elsewhere.

**Figure 6:**
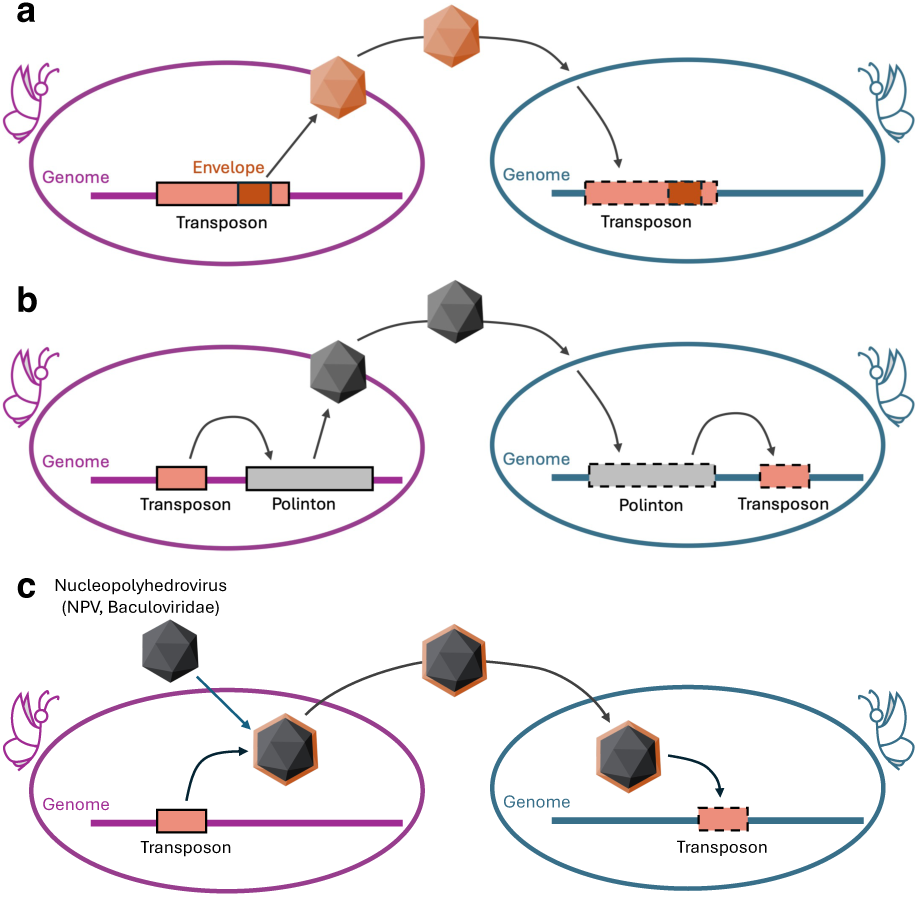
Hypothetical routes for horizontal transfer of TEs between insect genomes. **a,** A TE encoding an *envelope* gene enables the formation of infectious virus-like particles that move between hosts. **b,** A TE is carried as cargo within a polinton, which encodes capsid proteins, enabling particle formation and transfer between hosts. **c,** A TE integrates into a viral genome, which enables infection of multiple hosts and mediates TE transfer.

First, autonomous transfer is plausible for LTR retrotransposon MEFs shared between distant species that encode not only *gag* and *pol* but also *env* (envelope) proteins. The acquisition of *env*, likely from infectious retroviruses, enables receptor binding, membrane fusion, and formation of virus-like particles capable of infecting recipient cells [Malik et al., 2000]. Our example, the *Wallaby* element, is an *env* -encoding Bel/Pao LTR retrotransposon whose envelope retains the features of functional envelope proteins. By analogy, *env* -encoding elements of the Ty3/gypsy clade exhibit infectious properties and mediate intercellular transmission, including experimentally demonstrated horizontal transfer and infection of recipient germline cells—larval feeding of envelope-carrying gypsy/mdg4 particles generated novel germline insertions [Kim et al., 1994, Song et al., 1994, Pelisson et al., 2002]—and such endogenous retroviruses can infect neighbouring cells via virus-like particles [Senti et al., 2026]. While direct evidence for interspecies transfer remains limited, these findings support the potential of *env* containing TEs for autonomous horizontal transfer (Fig. 6a).

Second, when it comes to polintons as vectors for non-autonomous transfer, both moths and butterflies harbour active polintons that themselves show clear signatures of horizontal transfer, consistent with reports identifying Lepidoptera as an HT hub, although those studies did not discuss polintons [Reiss et al., 2019]. Previous evidence that polintons can cross species barriers supports host switching, but on deeper timescales than found here, and earlier work showed polintons can carry gene and TE cargo [Widen et al., 2023, Jeong et al., 2023]. Our results extend this by showing that polintons with TE insertions can remain mobile. We could not directly show a polinton carrying MEF cargo between hosts, but consider it plausible that broader sampling will reveal such cases. Together, polintons satisfy key prerequisites for acting as vectors of non-autonomous TE transfer (Fig. 6b).

Third, we detected active copies of MEFs shared among host species within a nucleopolyhedrovirus (baculovirus) infecting one of them. Given the well-documented bidirectional exchange of genetic material between viruses and host genomes, this is consistent with a role of viral infection in cross-species TE movement [Gilbert et al., 2016, 2014, 2010, Fraser et al., 1985, Muller et al., 2021] and points to baculoviruses as plausible vectors for at least some inferred HT events (Fig. 6c). The element, *piggyMITE*, is a non-autonomous *piggyBac*-like MITE found in two distant hosts and within a baculovirus able to infect at least one, raising the possibility that the virus ferried it between them. Equally, an earlier transfer by another agent followed by independent mobilization could have seeded the baculovirus insertions, so vector and direction remain provisional. Fittingly, *piggyBac* itself was first identified as an insertion in a baculovirus genome, so recovering a *piggyBac*-like MITE within HzNPV echoes the founding history of the superfamily [Cary et al., 1989, Wang et al., 1989].

Notably, several recurring HT hosts here—*Helicoverpa zea*, *Plutella xylostella*, *Pieris*, and *Aphis gossypii* —are agricultural pests against which baculoviruses such as HzNPV and HearNPV are mass-released as biopesticides [Sun, 2015], plausibly raising the viral exposure that could mediate TE transfer [Gilbert et al., 2014]. This suggests agroecosystems as a setting to test whether pest management accelerates mobilome exchange.

Beyond its primary scope, this study provides a collection of MEFs with direct evidence of activity from whole-genome alignments rather than inferred from copy number or similarity to known TEs, spanning 60 insect genomes and to be expanded in future work. Incomplete MEFs, comprising about 50% of candidates, were excluded pending further classification, highlighting a substantial and largely unexplored diversity that includes non-autonomous elements whose mobilization and emergence remain open questions.

Finally, the current design of Pannagram suggests several directions for future development. It is efficient in reference-free alignment because it avoids all-to-all comparison and uses a small set of references, running several-fold faster and with substantially lower peak memory than comparable tools at matched coverage and accuracy (see Benchmarking). Its early stages nonetheless use BLAST, whereas newer methods such as FastGA may offer speed improvements [Myers et al., 2025]; its modular alignment pipeline should facilitate such upgrades. Although Pannagram identifies MEFs, it does not yet classify them or annotate features such as LTRs, TIRs, and target site duplications, functions planned for a future Features module. More broadly, like current SV callers it struggles to resolve complex SVs with more than two allelic states; to address this we aim to incorporate ancestral recombination graph–based approaches to reconstruct the order of events in such loci.

## Acknowledgements

This work was supported in part by ERC Advanced Grant EPICLINES (to M.N.).

## Data availability

All genome assemblies analysed in this study are publicly available in GenBank; accession numbers are listed in Supplementary Table 1. The catalogue of mobile element families (MEFs) generated here, together with the horizontal-transfer candidate sequences, will be deposited in a public repository (Zenodo DOI to be added upon acceptance).

## Code availability

Pannagram is open source and available at https://iganna.github.io/pannagram/; the exact version used for this study will be archived with a citable DOI (Zenodo, to be added upon acceptance).

## Methods

### Multiple genome alignment

Pannagram is capable of constructing whole-genome alignments in three modes: preliminary, reference-based, and reference-free (Extended Data Fig. 9). In preliminary mode, all chromosomes from the genomes of interest are aligned to a reference genome. In this process, each chromosome is divided into windows of a specific length (1 kb by default). For each window, a BLAST search [Altschul et al., 1990] is performed to identify the best-matching location in the reference genome. Finally, during the linearization step, hits that deviate from the main linear trend are filtered out to ensure alignment accuracy. As a result, a multiple genome alignment of the most conserved syntenic regions is produced, which the user can explore through the generated correspondence plots. This mode is useful for examining the correspondence between chromosomes, for example, to verify that all chromosomes are in the same order across accessions. It also allows for the detection of large inversions, translocations, and rearrangements, as well as potential assembly issues that should be considered. Identifying such regions may help refine the assembly or make necessary adjustments and chromosomal rearrangements before performing the final full reference-free alignment.

The reference-based mode is an extension of the preliminary mode. It identifies all breaks in the preliminary alignment and attempts to align them with greater precision and accuracy using BLAST.

The reference-free mode builds upon the reference-based approach. It selects multiple accessions as references, either specified by the user or, by default, taken as the first accessions in alphabetical order, and performs a reference-based alignment for each. Then, it extracts only those positions that correspond to each other across all reference-based alignments, regardless of which reference was used. This results in highly syntenic reference-free blocks. Finally, the gaps between these blocks are stitched together using the original Pannagram pipeline, which combines abPOA, BLAST and MAFFT [Gao et al., 2021, Katoh et al., 2002].

### Chromotools

Relatively often, publicly available genomes of the same species do not follow the strict chromosome order, and the apparent large rearrangements, translocations, or inversions may represent either true biological differences or artefacts of genome assembly. Since Pannagram, in its fastest mode, aligns chromosomes sequentially (the first with the first, the second with the second, and so on), it is advisable to provide input genomes in which such rearrangements are restored to a consistent order.

Chromotools is a module that processes (cuts and joins) chromosomes to make all genomes of a species consistent with a single chosen reference genome. To use Chromotools, you first need to run Pannagram in the preliminary reference-based mode to build a skeleton alignment. Using this alignment, Chromotools module detects the borders of rearrangements and creates chimeric chromosomes to match the order of the reference genome. Chromotools can be used both within a single species and across different species.

### Simsearch

Pannagram contains a powerful and user-friendly module, Simsearch, to search nucleotide sequences by similarity and coverage across various contexts. Supported context types include (i) a set of sequences, (ii) a single genome, or (iii) a directory containing multiple genomes, covering the most typical and practically important use cases. Simsearch is a BLAST-based tool that post-processes and merges hits to meet the required similarity and coverage thresholds. The logic for handling sequence similarity and coverage depends on the context in which Simsearch is performed.

In the first context, when one set of sequences is compared to another set, Simsearch retains only those BLAST hits that meet the similarity threshold. It then evaluates coverage, which in simple cases is handled in a straightforward manner (Extended Data Fig. 10a–b). In more complex scenarios involving duplications and reverse complements, a more flexible coverage model is applied (Extended Data Fig. 10c–d). In particular, when multiple duplicated instances of one sequence collectively cover another sequence, the result is 100% coverage for both sequences (Extended Data Fig. 10c). The result of a Simsearch run is a table where each row represents a pair of sequences from the two sets that meet the similarity threshold and where at least one sequence satisfies the coverage threshold, the so-called nestedness condition.

When using Simsearch to query sequences in a genome or a directory with genomes, logic for handling sequence similarity and coverage is different. In simple cases, a sequence that meets the coverage and similarity thresholds can be found in the genome as a whole. In more complex cases, when the sequence aligns as two neighbouring BLAST hits in a genome (each characterized by its own similarity and length), Simsearch computes the cumulative similarity and two separate coverage scores for the entire region (Extended Data Fig. 10e). To confirm that the sequence was detected, the cumulative similarity and the sequence-side coverage must exceed their thresholds, while the genome-side coverage is enforced only optionally. The Simsearch output in the genomic context includes tables showing the copy number of each sequence per genome, as well as a GFF file annotating the genomic regions where these sequences were detected. Limitations or errors in detecting initial hits depend on BLAST.

### SNPs and Pi diversity

In Pannagram, SNPs are defined as all nucleotide positions in the whole-genome alignment that exhibit polymorphism between genomes, regardless of whether such positions are adjacent to each other or located within SVs. This inclusive definition reports all polymorphic sites without prior filtering, allowing users to apply custom filtering criteria as needed (e.g., retaining only polymorphisms without other polymorphisms within a ±2 nucleotide window).

The Features module of Pannagram outputs a VCF file containing SNPs, with the reference allele defined as the most abundant allele across the analysed genomes. The module also supports the calculation of nucleotide diversity (*π*) using VCFtools [Danecek et al., 2011] and PLINK [Chang et al., 2015].

### Structural Variants

There are different categories of events that can be classified as structural variants (SVs). In Pannagram, SVs are defined as loci in the whole-genome alignment where genomes demonstrate pres-ence–absence polymorphisms. Following this definition, SVs range in size from the simplest insertions or deletions of a single base pair to large genomic regions spanning multiple alternative alleles of varying lengths. In Pannagram, SVs are subdivided into simple and complex types, and simple SVs are characterized by the presence frequency, defined as the number of genomes carrying the presence allele in the pangenome locus.

Complex SVs often likely represent a combination of multiple events that are accumulated over time, that can make them difficult to interpret. In contrast, simple SVs usually result from single events and are therefore easier to analyse and more likely to reflect recent variation. For this reason, we recommend focusing downstream analyses on simple SVs.

SV-calling is performed using the module Features. In the output files, simple SVs are further subdivided into short (*<* 50 bp) and long (≥ 50 bp) SVs to facilitate their interpretation.

### Families of Mobile Elements

We hypothesize that a subset of SVs may result from the activity of mobile elements. To focus on active families of mobile element, we restrict our analysis to simple SVs. We define a mobile element family (MEF) as a group of SVs that exhibit nested sequence similarity, suggesting a common origin. Under this definition, a family is identified only if it is represented by at least three SVs across the analysed genomes. As a result, low-activity families that give rise to only a few SVs are not detected.

The pipeline for detecting MEFs follows a three-step logic. First, all sequences of SVs are compared against each other. Next, the skeleton (graph) of MEFs is constructed using highly represented sequences. Finally, the remaining SVs are added back to extend the graph of MEFs.

The detailed MEF-calling pipeline implemented in the Features module proceeds as follows. In the first step, only SVs that are at least 200 bp long are considered. These SVs are processed using the Simsearch module with default coverage and nested similarity cutoffs set to 85% (the 85–85 criterion). While the standard definition of MEFs uses an 80–80 criterion [Wicker et al., 2007], we use more stringent thresholds because in our experience this cutoff does not provide sufficient specificity. However, the user can specify alternative criteria.

In the second step, a graph of nestedness is constructed using the same 85% coverage cutoff (the similarity threshold is kept at 85%) and includes only SVs which exhibit mutual nestedness with other SVs (i.e. are represented at least twice in the dataset). Then some edges in this graph are filtered because they represent bypassed paths. For example, an edge connecting nodes 1 and 3 is removed if a path 1→2→3 exists. The resultant graph contains so-called “fork” and “umbrella” nodes: SVs that are nested within several other SVs, or SVs that contain several nested SVs, respectively. Edges associated with these nodes often connect SVs belonging to different MEFs. Therefore, outgoing edges from fork nodes and incoming edges to umbrella nodes are filtered. An edge is retained only if the shorter sequence aligns to either the beginning or the end of the longer one with at least 80% coverage, and the two nodes share at least 70% of their neighbouring edges. In other words, if a smaller sequence is contained within a larger one in the middle region, it is more likely to represent a parasitic family rather than a true relationship within the family. After this filtration, the graph is partitioned into families as connected components (the edges linking different families have already been removed by the shortcut, fork, and umbrella filtering above), yielding the skeleton of MEFs, though not yet in its final form.

In the third step, all nodes that were filtered out by the 90–90 criterion are added back. If a node can be connected to multiple MEFs, it is assigned to the MEF with which it shares at least 70% of its edges (the dominant effect). If a node is not connected to any of the skeleton MEFs, it initiates a new MEF. After this assignment, inter-community edges are removed once more, and the graph is re-partitioned into communities. The resultant graph is considered the graph of MEFs, in which each community represents a separate MEF.

For the manual extraction of Mobile Element Families, the Pannagram R library provides a set of user-friendly functions described in detail in the documentation. These functions are designed to be combined into a flexible, user-defined pipeline, allowing full control over the choice of parameters and filtering steps.

### Criteria for identifying SVs of autonomous TEs

We extracted all ORFs of at least 300 aa from RepBase TE sequences and all ORFs of at least 200 aa from SV sequences. Protein BLAST was then performed using SV-derived ORFs as queries against RepBase-derived ORFs, and only hits spanning more than 160 aa were retained. Each SV was assigned to the TE class corresponding to its longest BLAST hit. SVs classified in this way were considered proxies of autonomous TEs.

### Search for *piggyBac* autonomous element

We assessed whether candidate MEFs could be mobilized by autonomous *piggyBac* elements encoded either by viruses or by the host genomes of *H. zea* and *E. belemia*. We retrieved all *piggy-Bac* ORFs longer than 300 aa from Repbase, yielding 572 candidate transposases, and used them as queries in tBLASTn searches against available *Helicoverpa zea nucleopolyhedrovirus* (HzNPV) genomes (KM596835.1, AF334030.1, KJ004000.1). No matches longer than 124 aa were detected, well below the minimal length reported for functional *piggyBac* transposases (*>*300 aa) [Lyu et al., 2022].

Second, we performed tBLASTn searches against two available *E. belemia* genomes (GCA 964340465.1 and GCA 964340475.1) and two *H. zea* genomes (GCA 022343045.1 and GCA 022581195.1) using the same approach. This identified 4,919 loci in *E. belemia* and 3,422 loci in *H. zea* containing regions with similarity to *piggyBac* transposases over lengths exceeding 300 aa. For each hit, we extracted 3 kb of flanking sequence on both sides and screened for candidate terminal inverted repeats (TIRs), defined as 13 bp reverse-complement motifs allowing up to one mismatch.

We identified 296,910 candidate TIR pairs, none of which were identical to the TIRs of our candidate element. The closest match was detected in *H. zea* and differed by a single nucleotide, whereas the second closest, from *E. belemia*, differed by three nucleotides (Extended Data Fig. 8). These results do not support the presence of a canonical autonomous element in the analysed genomes, although unidentified *piggyBac* elements with relaxed TIR specificity may still provide transposase activity.

### Controls for cross-species contamination

Because near-identical sequences shared between species can in principle arise from sample cross-contamination or from host DNA carried over into microbial and viral assemblies, we required each candidate horizontal-transfer case to satisfy controls that free contaminant DNA cannot produce. First, we retained only elements with clean insertion boundaries, that is, flanked by terminal inverted or direct repeats and target-site duplications in the assembled genome. Second, we verified that a shared element occupied distinct insertion sites, with different host flanking sequence, in each species, which is expected for genuine genomic integration but not for a contaminating fragment. Third, we required the shared copies to differ by SNPs and indels rather than to be identical, since contamination would yield near-perfect identity. For the baculovirus case, we further required the shared region to be present in all independent HzNPV assemblies from different isolates, so that a single contaminated assembly could not account for the signal.

## Extended Data Figures

**Extended Data Fig. 1:**
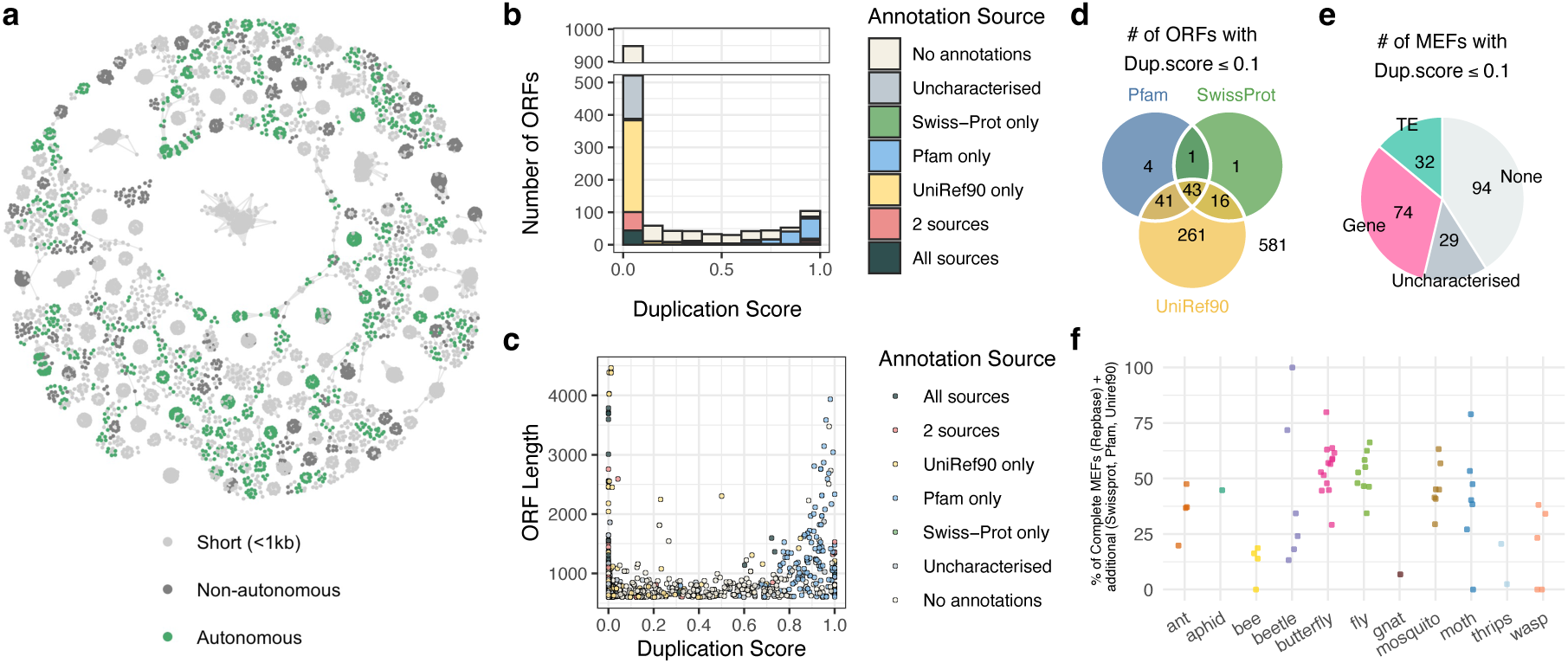
Annotation of long ORFs in incomplete MEFs. **a,** Graph-based representation of MEFs in *Anopheles arabiensis* colored by autonomous elements. **b,** Distribution of annotation sources for ORFs from incomplete MEFs from all insects across duplication-score. ORFs longer than 200 aa were analysed. Duplication score was defined as the fraction of 11-mers occurring at least twice within the nucleotide sequence of ORFs. ORFs were annotated using DIAMOND [Buchfink et al., 2021] searches against Swiss-Prot and UniRef90, and HMMER [Eddy, 2011] searches against Pfam. Uncharacterized proteins and DUFs were excluded. Pfam-only annotations were enriched at high duplication scores, suggesting lower annotation confidence, whereas ORFs supported by all three methods were concentrated in the lowest duplication-score bin (≤0.1). **c,** Distribution of ORF lengths across duplication-score bins. Long ORFs, likely corresponding to TE-related sequences, were concentrated in the lowest duplication-score bin (≤0.1). **d,** Congruence between annotation methods within the lowest duplication-score bin. Almost all ORFs annotated by Swiss-Prot or Pfam were consistently supported by at least one other method. **e,** ORFs were mapped to MEFs and classified as TE-related, gene-coding, or “Uncharacterized” (uncharacterized proteins and DUFs). The latter category may include novel TEs or unknown mobilized genes. **f,** The proportions of autonomous MEFs and MEFs containing TE-related ORFs remained unchanged after inclusion of the latter.

**Extended Data Fig. 2:**
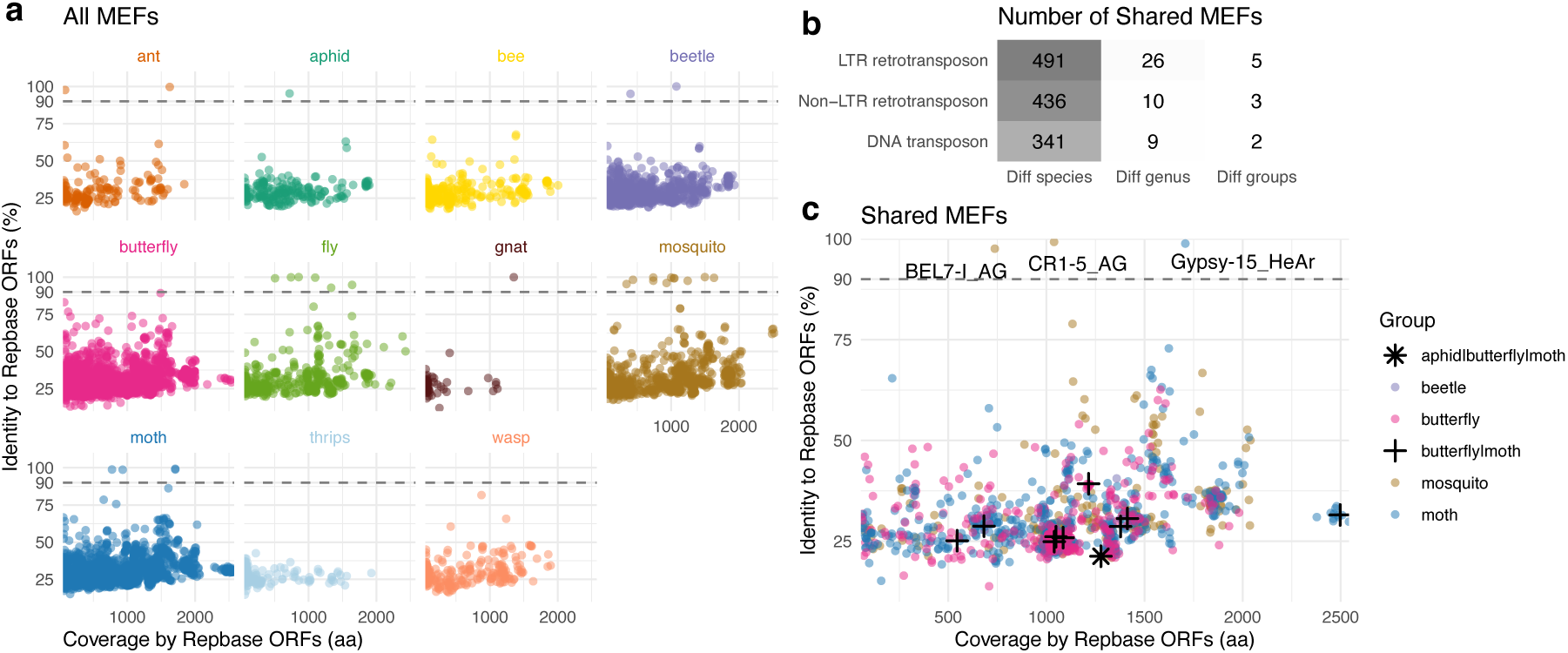
Classification of MEFs. **a,** Similarity of MEF ORFs to ORFs in Repbase. **b,** Number of MEFs from different TE classes shared between different genus and higher taxonomic groups. **c,** Similarity of MEF ORFs to ORFs in Repbase, shown only for MEFs shared between species.

**Extended Data Fig. 3:**
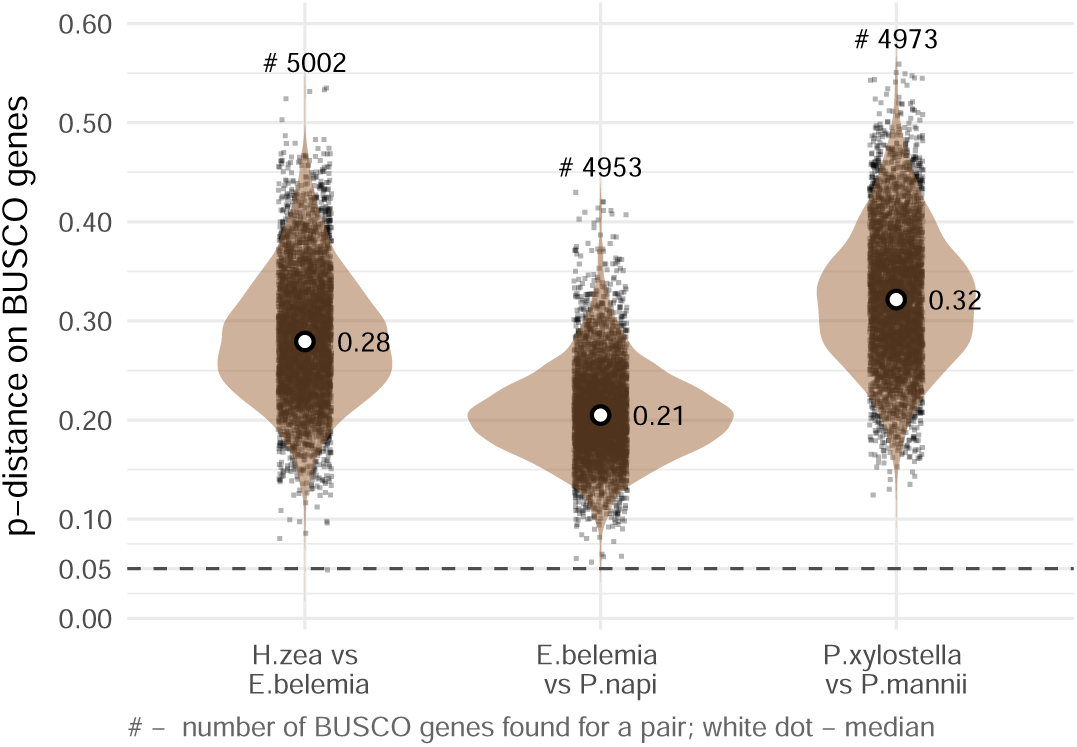
Genome-wide divergence between lepidopteran species pairs estimated from BUSCO genes. For each species pair, complete single-copy BUSCO orthologues from the lepidoptera odb10 lineage were aligned and compared. Pairwise p-distance was calculated for each shared gene as the proportion of mismatched positions across the aligned length. Violin plots show the distribution of per-gene p-distances. White dots indicate medians, and # gives the number of shared BUSCO genes in each comparison. The dashed horizontal line marks *p* = 0.05 as a visual reference. This 5% divergence corresponds to the 95% identity threshold used to call MEFs shared across species: because every host pair shown is far more diverged than this line, the shared MEFs are much more similar than expected from host divergence alone, consistent with horizontal transfer.

**Extended Data Fig. 4:**
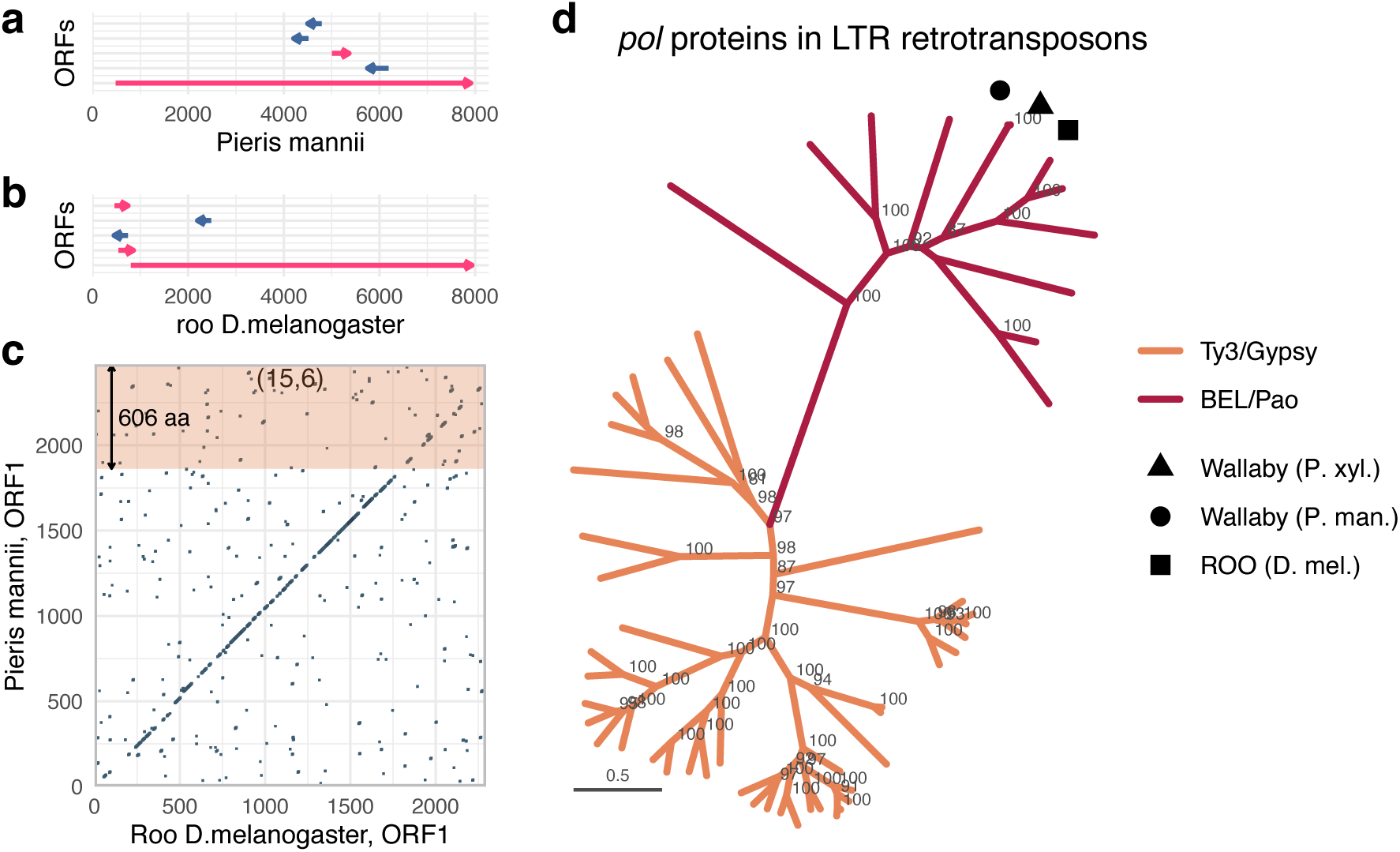
Horizontal transfer of TEs. **a,** Predicted open reading frame (ORF) organization of the *P. mannii* mobile element. Arrows indicate the position and orientation of individual ORFs along the full-length sequence. **b,** ORF organization of the *roo* element from *D. melanogaster*. **c,** Dot plot of the amino acid sequence of the ORF from the representative *P. mannii Wallaby* element and the ORF from the *D. melanogaster roo* element, both comprising *gag*, *pol* and *env* protein regions. The region corresponding to the *env* protein in *P. mannii* is highlighted in orange. **d,** Phylogenetic tree of the *pol* gene from the representative *Wallaby* MEF sequence and well-characterized LTR retrotransposons, placing *Wallaby* in a distinct branch of the Belpaoviridae (Bel/Pao) family, distant from the *roo* element of *D. melanogaster*.

**Extended Data Fig. 5:**
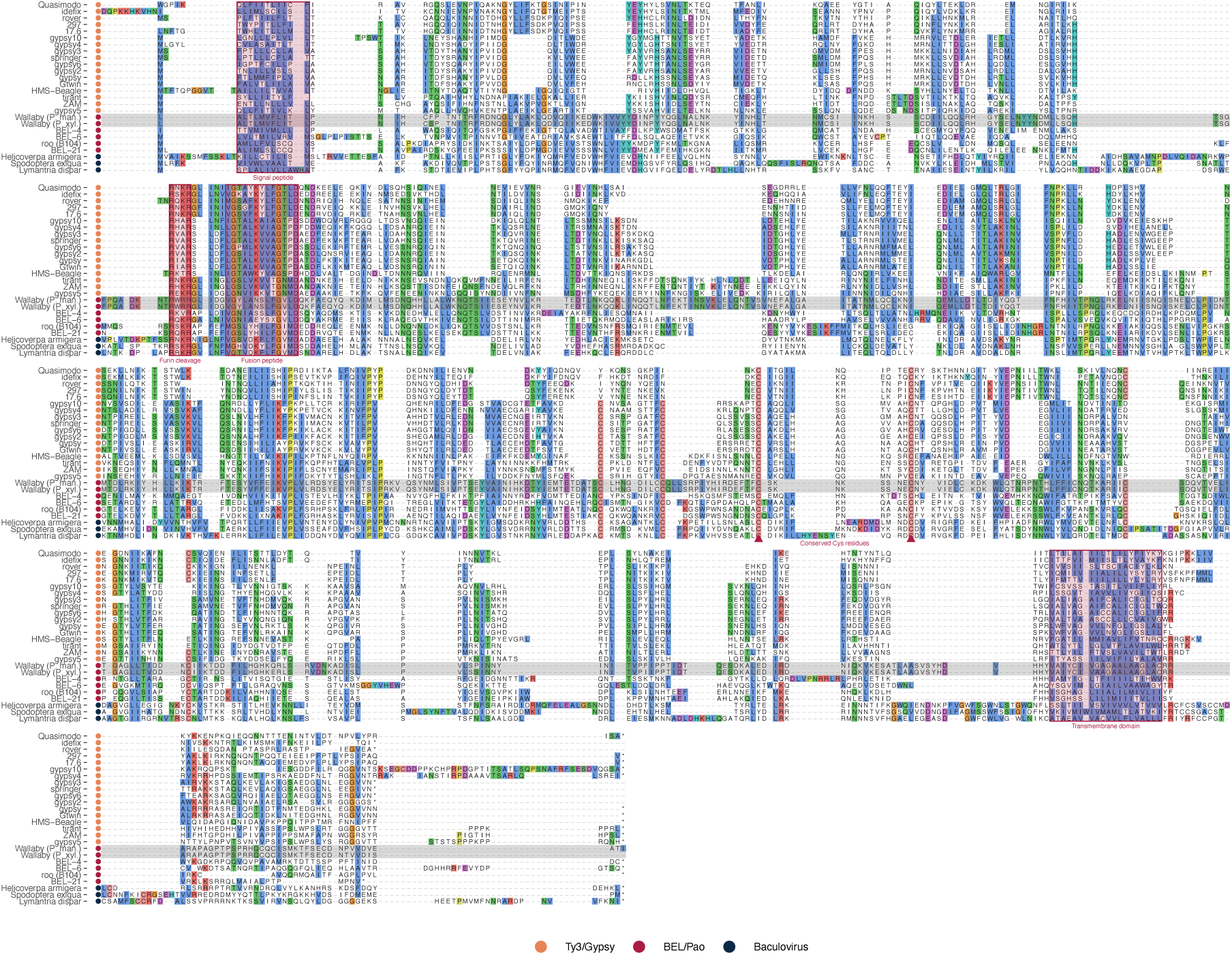
*Wallaby* MEFs encode Env proteins that retain the conserved architecture of functional envelope proteins. Multiple sequence alignment of Env proteins encoded by *Wallaby* MEFs, insect Ty3/Gypsy errantiviruses, BEL/Pao LTR retrotransposons, and baculovirus sequences (outgroup). *Wallaby* MEF sequences are shaded in grey. Red annotations mark the conserved Env-like features: the N-terminal signal peptide, furin cleavage site, fusion peptide, conserved cysteine residues putatively involved in disulfide-bond formation, and the transmembrane domain.

**Extended Data Fig. 6:**
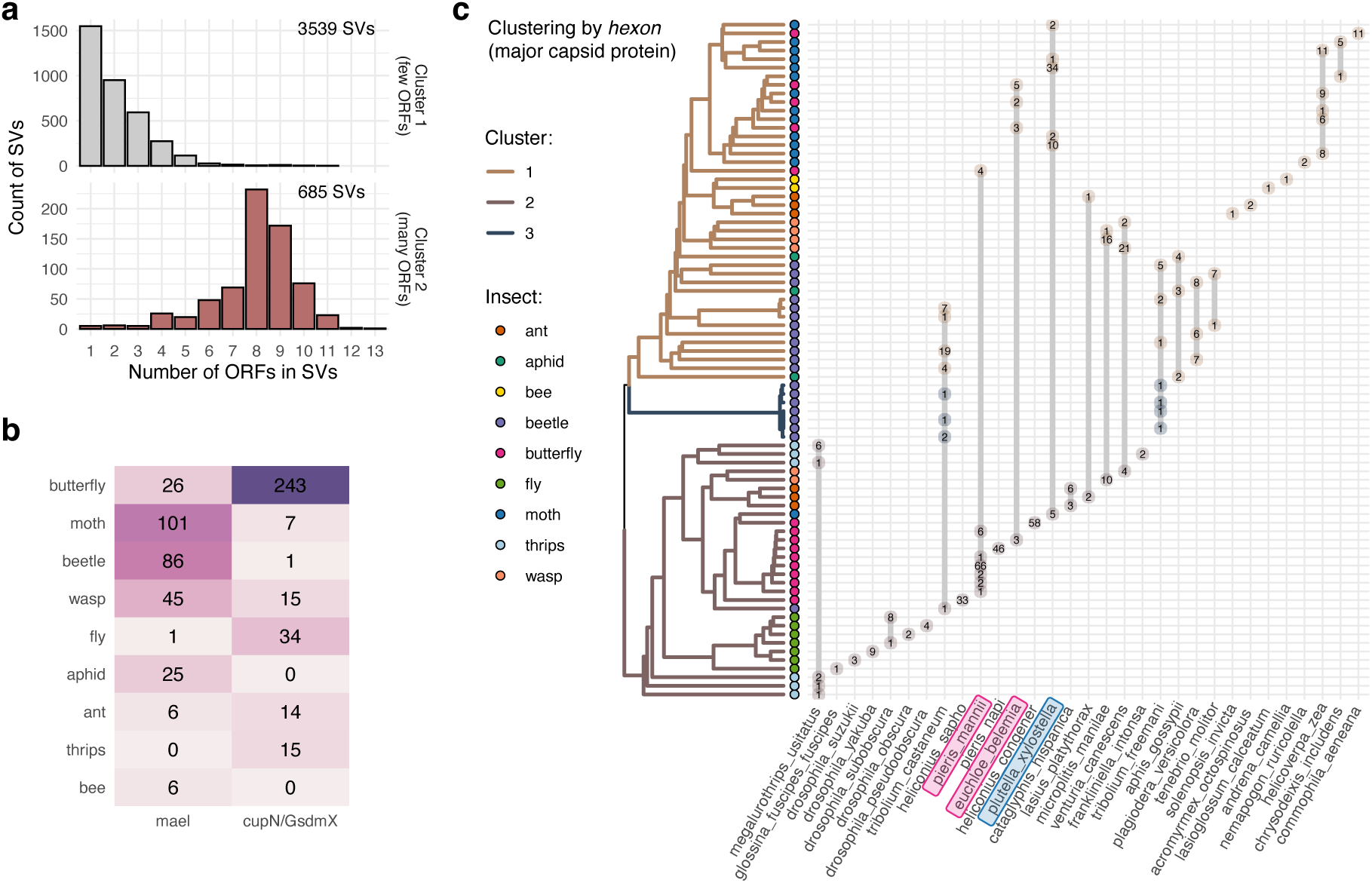
Horizontal transfer of polintons. **a,** Distribution of the number of ORFs (≥150 aa) per SV in the two clusters (”Few ORFs” and “Many ORFs”). **b,** Number of polinton-like SVs from the two groups ( *mael* -associated and *cupiennin/gasderminX* -associated) across different insect groups. **c,** Clustering of *hexon* proteins from polinton-like SVs. Numbers in circles indicate the number of SVs in species.

**Extended Data Fig. 7:**
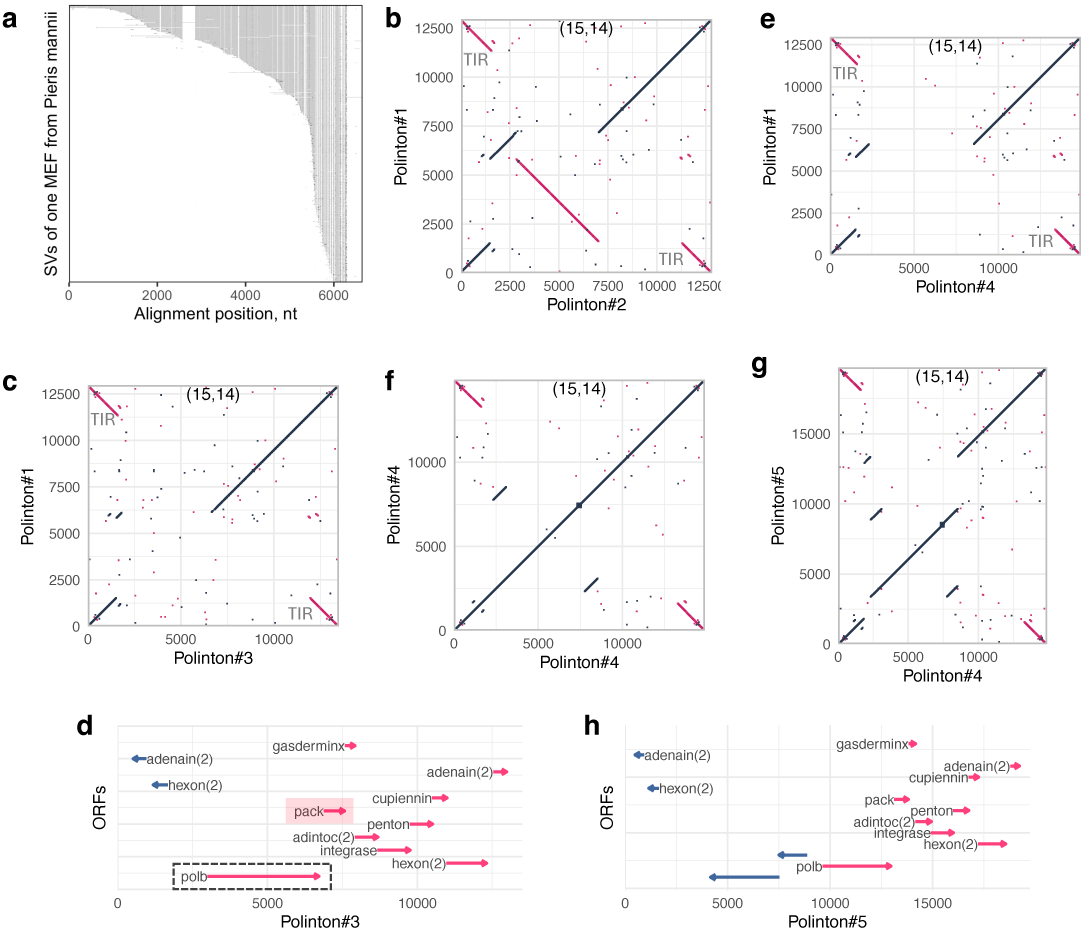
TE cargo in polintons. **a,** Multiple sequence alignment of SVs forming a single LINE-associated MEF in *Pieris mannii* (the cargo of the polinton-related SVs). Each row is an individual SV, sorted by length; grey indicates identity to the top (longest) sequence, black indicates mismatches, and white indicates gaps. The arrow labelled “SV” marks a variant within the MEF that is also observed in polinton elements. **b,** Dot-plot comparison of the two cargo-carrying polinton-related SVs, Polinton#1 vs Polinton#2 (window size 15; minimum matches 14). **c,** Dot-plot comparison of Polinton#1 vs Polinton#3 (window size 15; minimum matches 14). **d,** Gene organization of Polinton#3, which retains the *polB* gene. **e,** Dot-plot comparison of Polinton#1 vs Polinton#4 (window size 15; minimum matches 14). **f,** Example of a polinton-related SV with an insertion of a TE (LTR retrotransposon). **g,** Another polinton-related SV carrying the same TE insertion, together with an additional rearrangement, indicating a common origin of the two SVs and continued activity after TE insertion. **h,** ORF organization of the polinton-related SV shown in subplot **f**. The two blue-coloured ORFs between 4 kb and 8 kb correspond to ORFs in the inserted TE.

**Extended Data Fig. 8:**
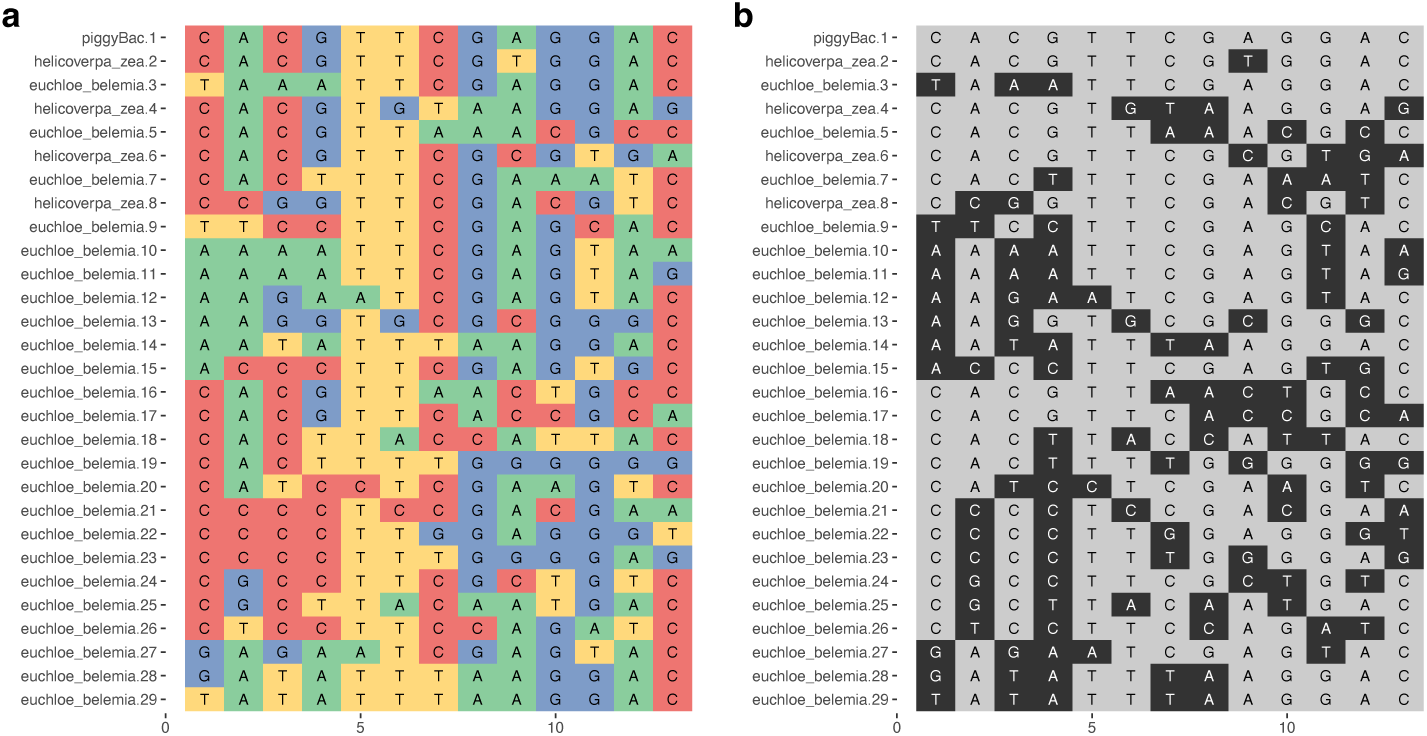
TIR motifs around *piggyBac* transposases. **a**, Alignment of motifs differing from the target motif by no more than five substitutions. The first line shows the target motif; other lines are labelled by organism and line number. **b**, The same alignment coloured by differences relative to the target motif. Grey indicates identical nucleotides, black - different.

**Extended Data Fig. 9:**
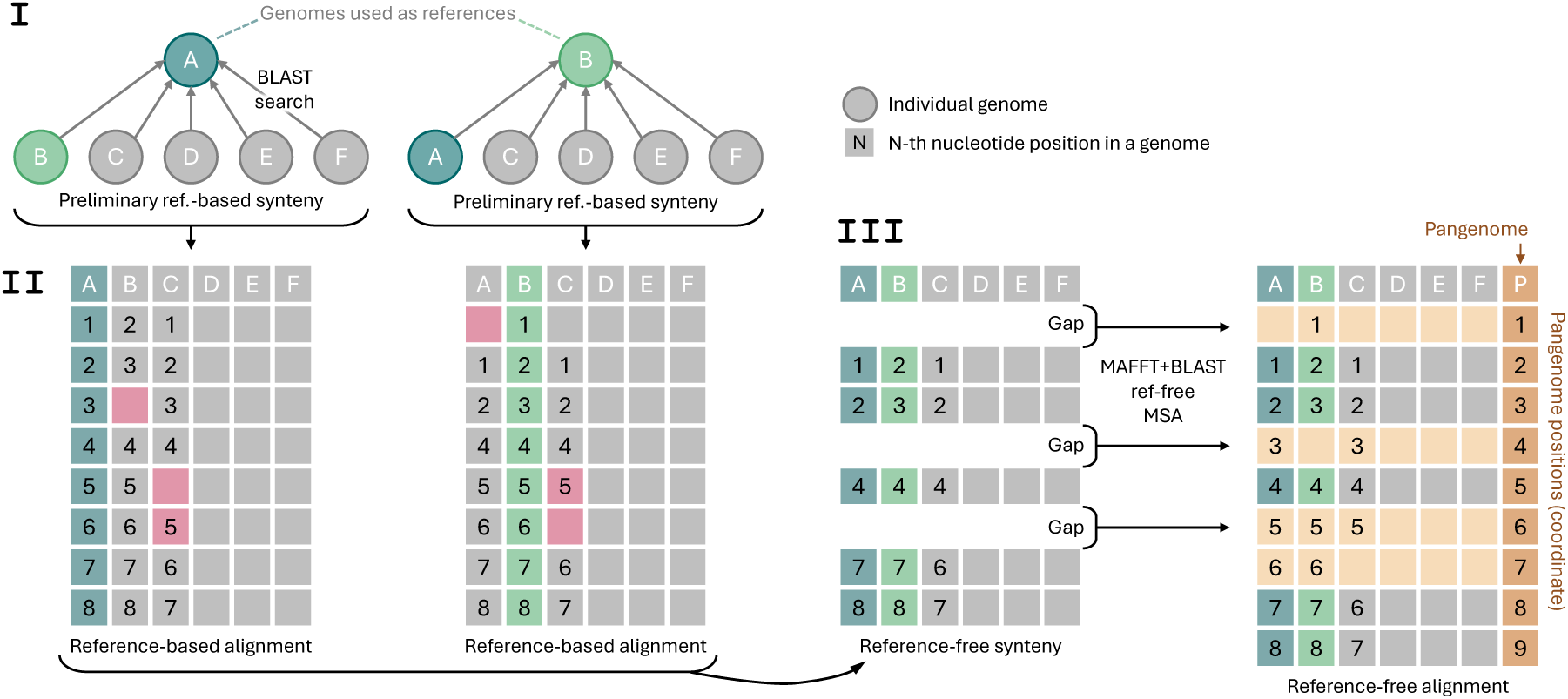
Scheme of reference-free alignment in Pannagram. Stage **I**: preliminary reference-based synteny is constructed using a selected set of reference genomes (chosen either randomly or deliberately). This step provides an initial skeleton of the alignment and identifies preliminary syntenic blocks. Stage **II**: reference-based alignments are built for each chosen reference, representing correspondences between nucleotide positions in all genomes as matrices of matched positions. Stage **III**: only highly syntenic positions that are consistent across different reference choices are retained to define reference-free synteny blocks. The gaps between these blocks are then filled using heuristics based on BLAST and MAFFT, resulting in the final reference-free alignment and pangenome coordinate system.

**Extended Data Fig. 10:**
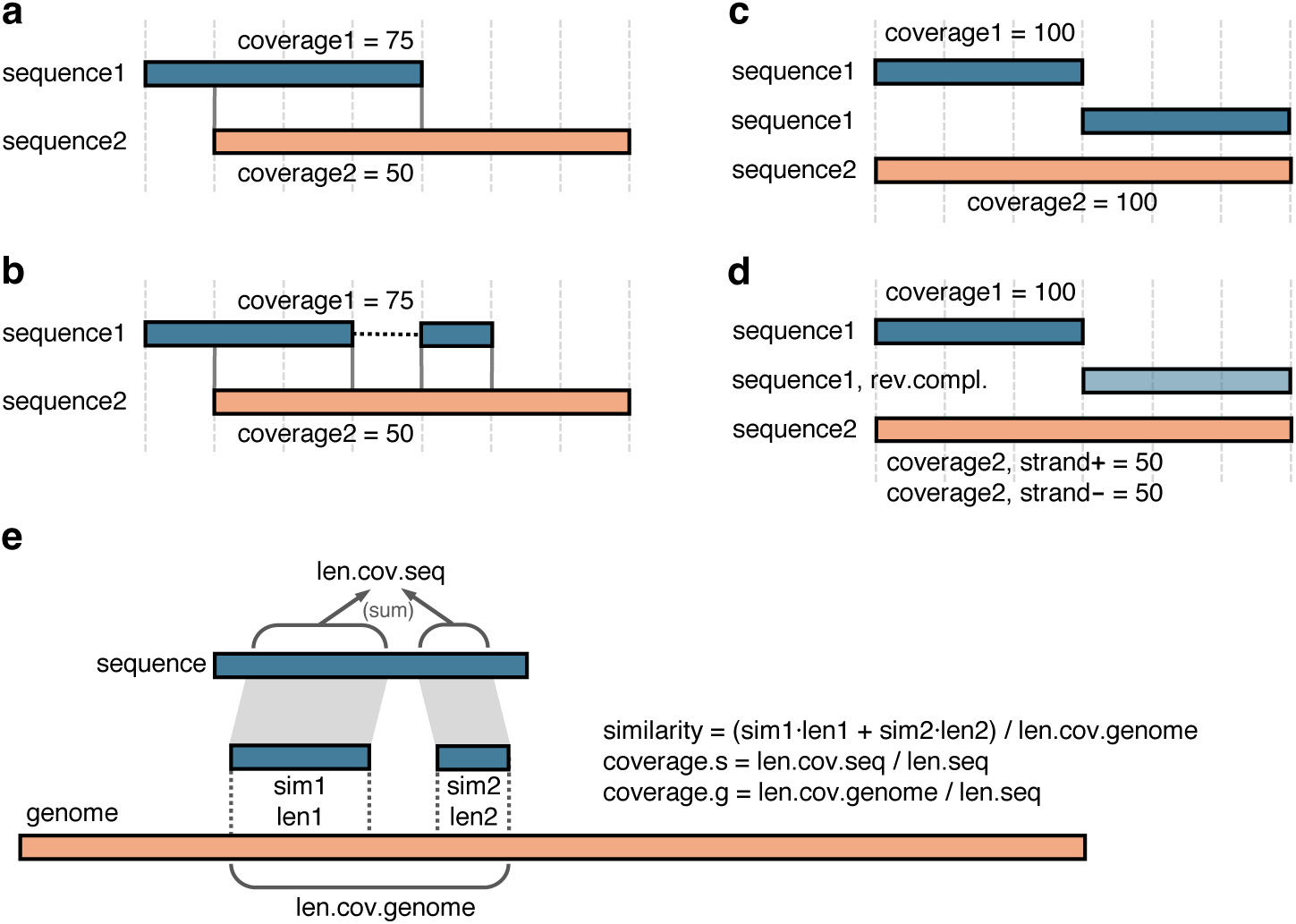
Coverage calculation in different alignment scenarios in the Simsearch module. **a**, Two partially overlapping sequences of different lengths. **b**, One sequence covers another with an internal gap. **c**, One sequence is duplicated relative to the other. **d**, One sequence is duplicated with an inversion relative to the other. **e**, Definition of the coverage variables when one sequence is mapped to a genome.

