## Supplementary Information for "Pannagram: unbiased pangenome alignment and mobilome calling across insects extends horizontal transfer from transposons to polintons"

Pannagram: reference-free pangenome alignment and mobilome calling

This Supplementary Information reports the full benchmarking of pannagram underlying the Benchmarking section of the main text: (i) computational cost and scaling against reference-free multiple-genome aligners (Supplementary Note S1); (ii) alignment accuracy on simulated genomes with known homology (Supplementary Note S2); (iii) the reference-based (`-ref`) mode against pairwise whole-genome aligners, with the word-size parameter sweep (Supplementary Note S3); (iv) pangenome mobile-element families against de-novo TE annotation with EDTA (Supplementary Note S4); and (v) the Simsearch module against RepeatMasker.

### S1 Computational benchmark against reference-free multiple-genome aligners

#### S1.1 Aim and scope

This benchmark positions pannagram against established *reference-free multiple whole-genome* aligners on computational cost and its scaling with the number of genomes. Pairwise aligners (e.g. minimap2, MUMmer) form a different methodological class and are not included here. The three comparators are Progressive Cactus, progressiveMauve and SibeliaZ. Because these tools emit different products (Table S1), runtime is interpreted alongside what each tool actually computes rather than in isolation.

Table S1: Output class of each aligner. Runtimes are only strictly comparable within the same output class.

| Tool | Reference-free | Primary output |
|---|---|---|
| pannagram | yes | base-level MSA + per-site SNP/SV calls |
| Progressive Cactus | yes | base-level MSA + ancestral sequences (HAL) |
| progressiveMauve | yes | locally-collinear blocks, base-level (XMFA) |
| SibeliaZ ($-n$) | yes | synteny-block coordinates (no base-level) |

#### S1.2 Datasets

Two intraspecific panels of chromosome-scale assemblies were used. For each panel a fixed, ordered pool of accessions was defined and the first $N$ accessions were taken at scaling point $N$; all tools received the identical FASTA set at each point.

- *Drosophila melanogaster*: six chromosome-scale sequences per assembly, ≈134.5–134.7 Mb per genome; $N \in \{2, 5, 8, 11\}$.
- *Anopheles coluzzii*: five chromosome-scale sequences per assembly, ≈246.8–249.5 Mb per genome; $N \in \{2, 5, 8\}$.

Input identity across tools was verified by recording an MD5 checksum of every genome supplied to every run. Both panels are intraspecific and of low divergence: pairwise Mash distances (sketch size $10^4$, $k$=21) were 0.0046–0.0064 (mean 0.0054; ≈99.4–99.5% ANI) for *D. melanogaster* and 0.0056–0.0203 (mean 0.0150; ≈98.0–99.4% ANI) for *A. coluzzii*.

#### S1.3 Hardware and scheduling

All runs were executed on a single cluster partition of dual-socket Intel Xeon Platinum 8368 @ 2.40 GHz nodes (76 physical cores, 1 TB RAM). Each (tool, scaling-point) combination was submitted as one SLURM job with `--cpus-per-task=16 --mem=256G --time=24:00:00`. Elapsed wall-clock time and maximum resident set size (peak RSS) were recorded with `/usr/bin/time -v`; peak RSS in kilobytes was converted to gigabytes.

#### S1.4 Software versions and commands

- **pannagram** v0.1.0 (commit 5da3fc3), reference-free MSA:
  pannagram -path_in <dir> -path_out <proj> -nchr <NCHR> -cores 16
  (defaults: part_len 1000, word_size 28, word_size_gap 15).

- **Progressive Cactus** binary release v3.2.1:
  cactus <jobstore> <seqFile> out.hal --maxCores 16; guide tree a balanced caterpillar with uniform 0.01 branch lengths (same species, topology ≈ irrelevant); FASTA headers truncated at the first whitespace as required by Cactus.

- **progressiveMauve** build 2015-02-13:
  progressiveMauve --output=out.xmfa g1 ...gN (defaults).

- **SibeliaZ** v1.2.7 (TwoPaCo 1.1.0, spoa 4.1.5):
  sibeliaz -n -t 16 -o <out> all.fa — run in block mode ($-n$); the base-level (spoa) step was not run because it does not scale to genomes of this size.

#### S1.5 Metrics and reproducibility of the timings

Primary metrics are wall-clock time (s) and peak memory (GB). CPU utilisation (% of one core, from /usr/bin/time) is reported as a secondary measure of parallel efficiency. The scaling points in Table S3 are single runs; to establish that single runs are representative, every tool was additionally run in triplicate at the two-genome point of both panels (Table S2). Run-to-run variation is small (coefficient of variation 1.6–4.0%) for seven of the eight tool×species combinations, so single-run timings are reliable at the reported precision. The one exception (pannagram on *Anopheles*, CV 20%) is driven by a single slow replicate (1,613 s vs 1,050/1,111 s) attributable to co-tenancy on the shared node; the median is used.

Table S2: Triplicate timings at $N$=2 (seconds): median, standard deviation and coefficient of variation.

| Species | Tool | Median (s) | SD (s) | CV (%) |
|---|---|---|---|---|
| *D. melanogaster* | pannagram | 387 | 9 | 2.4 |
| | SibeliaZ | 49 | 1 | 2.5 |
| | Cactus | 652 | 15 | 2.4 |
| | Mauve | 1,039 | 17 | 1.6 |
| *A. coluzzii* | pannagram | 1,111 | 252 | 20.1 |
| | SibeliaZ | 167 | 3 | 1.9 |
| | Cactus | 1,131 | 43 | 3.7 |
| | Mauve | 1,509 | 61 | 4.0 |

#### S1.6 Results

Table S3 reports wall-clock time, peak memory and CPU utilisation for all four tools at every scaling point.

Table S3: Wall-clock time, peak memory and CPU utilisation for reference-free multiple-genome alignment as a function of the number of genomes $N$. All runs completed; the slowest (Mauve, eight *Anopheles* genomes) took 21.0 h.

| Species | Tool | $N$ | Time (s) | Peak RAM (GB) | CPU (%) |
|---|---|---|---|---|---|
| *D. melanogaster* | pannagram | 2 | 384 | 3.6 | 118 |
| | pannagram | 5 | 1,221 | 5.8 | 110 |
| | pannagram | 8 | 2,219 | 8.2 | 109 |
| | pannagram | 11 | 3,079 | 9.7 | 106 |
| | SibeliaZ | 2 | 48 | 2.8 | 1089 |
| | SibeliaZ | 5 | 91 | 2.8 | 1137 |
| | SibeliaZ | 8 | 144 | 3.8 | 1213 |
| | SibeliaZ | 11 | 198 | 5.2 | 1211 |
| | Cactus | 2 | 651 | 24.5 | 224 |
| | Cactus | 5 | 2,569 | 26.9 | 589 |
| | Cactus | 8 | 5,607 | 27.9 | 729 |
| | Cactus | 11 | 9,093 | 26.9 | 783 |
| | Mauve | 2 | 1,024 | 6.6 | 99 |
| | Mauve | 5 | 7,858 | 16.8 | 99 |
| | Mauve | 8 | 22,877 | 24.0 | 99 |
| | Mauve | 11 | 38,797 | 31.3 | 99 |
| *A. coluzzii* | pannagram | 2 | 1,111 | 7.0 | 103 |
| | pannagram | 5 | 4,589 | 11.5 | 95 |
| | pannagram | 8 | 7,745 | 16.0 | 100 |
| | SibeliaZ | 2 | 161 | 3.1 | 1133 |
| | SibeliaZ | 5 | 307 | 8.7 | 1093 |
| | SibeliaZ | 8 | 430 | 13.6 | 1044 |
| | Cactus | 2 | 1,131 | 33.0 | 556 |
| | Cactus | 5 | 5,606 | 46.8 | 984 |
| | Cactus | 8 | 11,484 | 43.0 | 1015 |
| | Mauve | 2 | 1,509 | 8.6 | 99 |
| | Mauve | 5 | 20,224 | 22.9 | 99 |
| | Mauve | 8 | 75,514 | 44.1 | 99 |

#### S1.7 Observations

Peak memory is the clearest differentiator. pannagram used 3.6–9.7 GB (*Drosophila*) and 7.0–16.0 GB (*Anopheles*), an order of magnitude below Cactus (24.5–46.8 GB) and, at scale, below Mauve (up to 31.3 GB), while still producing a base-level alignment with per-site variant calls. pannagram's runtime scaled approximately linearly with $N$ and was 3–13× faster than Cactus and Mauve at the largest panels; Mauve scaled super-linearly in both time and memory, needing 21.0 h and 44 GB for the eight-genome *Anopheles* panel (versus 2.2 h and 16 GB for pannagram). SibeliaZ was fastest and lightest, but only when run in block mode (no base-level alignment); its throughput reflects a $k$-mer / de-Bruijn-graph synteny-detection strategy rather than progressive base-level alignment. CPU utilisation shows that SibeliaZ is heavily multithreaded (∼11×), Cactus parallelises increasingly with $N$, Mauve is effectively single-threaded (99%), and pannagram is largely serial (∼1.1×)—so pannagram's speed derives from its algorithm rather than from using the 16 available cores, leaving headroom for future parallelisation.

#### S1.8 Alignment coverage

Runtime is only interpretable alongside how much sequence each tool actually aligns. Table S4 reports, at the five-genome point, the fraction of each genome placed in an alignment column with at least one other genome, averaged over the five genomes (computed from the retained alignment outputs: pannagram's pangenome HDF5, Cactus HAL→MAF, Mauve XMFA, SibeliaZ block GFF).

Table S4: Fraction of each genome aligned to ≥1 other genome ($N$=5, mean of five genomes). SibeliaZ is block-level: its value is synteny-block coverage, an upper bound that is not base-level and therefore not directly comparable.

| Species | pannagram | Cactus | Mauve | SibeliaZ* |
|---|---|---|---|---|
| *D. melanogaster* | 0.938 | 0.933 | 0.847 | 0.983* |
| *A. coluzzii* | 0.807 | 0.864 | 0.789 | 0.967* |

pannagram's base-level coverage matches Cactus on *Drosophila* (0.938 vs 0.933) and is somewhat lower on the more divergent, more repetitive *Anopheles* panel (0.807 vs 0.864); progressiveMauve aligns the least on both (0.79–0.85), consistent with its poor structural-variant accuracy (Supplementary Note S2). SibeliaZ's high value reflects permissive block-level synteny rather than base-level homology.

#### S1.9 Why pannagram is cheaper than Cactus and Mauve

The runtime and memory gap is algorithmic—pannagram performs strictly less computation. Progressive Cactus computes, per run, all-pairs LASTZ alignments along a guide tree, a cactus graph encoding rearrangements, base-level refinement, *and* reconstructed ancestral sequences at every internal node, orchestrated through the Toil engine with an HDF5 (HAL) assembly. progressiveMauve computes anchored locally-collinear blocks with an iterative refinement that scales super-linearly in genome number. pannagram, by contrast, seeds synteny by BLAST against the reference chromosomes and applies expensive MSA (MAFFT) only to the short residual gap loci between anchors; it reconstructs no ancestral genomes and builds no rearrangement graph. On intraspecific panels (≈99% identity) almost the whole genome is collinear, so the anchor-and- fill strategy leaves little work for the costly step, whereas Cactus and Mauve run their full machinery regardless of divergence. The CPU-utilisation figures make this concrete: pannagram overtakes Cactus while using ∼1 core on average versus Cactus's 7–10, so its advantage is lower total work, not better parallelism—and it holds the base-level alignment in far less memory because it never materialises the graph or the ancestral sequences. The trade-off is that Cactus's extra cost buys capabilities pannagram does not provide (ancestral genomes; an explicit rearrangement/duplication graph); for reference-free pangenome SNP/SV extraction within a species, that machinery is not needed.

#### S1.10 Limitations

(i) Scaling points are single runs (triplicates only at $N$=2, Table S2); jobs shared multi-tenant nodes, so peak memory is robust but wall-clock times carry scheduler / co-tenancy noise. (ii) SibeliaZ was run in block mode only and is therefore not directly comparable on the time axis to the base-level aligners. (iii) Both panels are intraspecific (low divergence), a regime that favours $k$-mer-based synteny detection. (iv) A thread-scaling sweep and a mammalian-scale genome were not run. (v) This benchmark measures computational cost and coverage only; alignment accuracy against a ground truth is assessed separately (Supplementary Note S2).

### S2 Alignment accuracy on simulated genomes

#### S2.1 Rationale and metric

Computational cost (Supplementary Note S1) is only meaningful alongside accuracy. Because the four tools emit different products, we score them on the one thing they all produce—a base-level alignment—rather than on tool-specific variant calls. From a simulation with a *known* ancestor and descendants we know exactly which ancestor base is homologous to which descendant base; each tool's alignment is compared against this truth. For every tool we extract, from its alignment output only (for pannagram, its position correspondence matrix; *not* its `features` variant caller), the set of ancestor→descendant aligned base pairs and compute

$$\text{recall} = \frac{\text{correctly aligned homologous pairs}}{\text{all true homologous pairs}}, \quad \text{precision} = \frac{\text{correctly aligned}}{\text{all pairs the tool proposes}},$$

and their harmonic mean $F_1$.

#### S2.2 Simulation and scoring

Ground truth was generated with simuG. To keep every truth call in the ancestor coordinate frame without composing multiple coordinate maps, SNPs and structural variants were placed in *separate* datasets, each descendant carrying a single edit class: (i) a **SNP** dataset (SNPs + short indels) and (ii) an **inversion** dataset (inversions 5–50 kb; the classic base-level-aligner discriminator—CNV duplications produce one-to-many homology that a pairwise base-level metric cannot score and were excluded). Each dataset was produced at two divergence levels ("low", "high"), four descendants each, with fixed seeds. The ancestor was included as a genome in every alignment so that all tools could be projected onto ancestor coordinates. Per-tool aligned pairs were obtained from: pannagram's HDF5 correspondence matrix; Cactus `hal2maf` → MAF; progressiveMauve XMFA; each parsed into the same ancestor-indexed representation and scored identically. Truth reconstruction was validated by exact reconstruction of descendant length.

#### S2.3 Results: *E. coli* (4.6 Mb)

Table S5 gives the mean over four descendants. SibeliaZ is absent: its base-level (spoa) alignment could not run at this scale (out-of-memory even at 256 GB), so it produces synteny blocks only and cannot be scored on base-level homology.

#### S2.4 Results: *Drosophila* (32 Mb chromosome)

Table S6 repeats the experiment on the largest *D. melanogaster* chromosome (Chr5, 32.3 Mb), an insect-scale, eukaryotic single-chromosome test.

At eukaryotic chromosome scale the picture is consistent with, and sharper than, *E. coli.* pannagram is robust everywhere ($F_1 \geq 0.997$) and is the best or tied-best method on inversions at both divergence levels. Cactus is marginally ahead on pure SNPs (recall ≈1.0 vs pannagram's 0.996) but pannagram matches it on structural variants at a fraction of the runtime and memory (Supplementary Note S1). progressiveMauve degrades markedly at this scale—its recall falls to ≈0.86 even on SNPs and to 0.72–0.80 on inversions—so it is both the slowest and the least accurate base-level method here.

Table S5: Alignment accuracy on simulated *E. coli* (mean of four descendants). "low"/"high" are ≈0.5%/2% SNP divergence for the SNP sets and 25/50 inversions per descendant for the inversion sets.

| Variant / divergence | Tool | Recall | Precision | $F_1$ |
|---|---|---|---|---|
| SNP, low | pannagram | 0.9994 | 0.9998 | 0.9996 |
| | Cactus | 0.9998 | 0.9998 | 0.9998 |
| | Mauve | 0.9995 | 0.9998 | 0.9996 |
| SNP, high | pannagram | 0.9979 | 0.9991 | 0.9985 |
| | Cactus | 0.9991 | 0.9991 | 0.9991 |
| | Mauve | 0.9983 | 0.9991 | 0.9987 |
| Inversions, low | **pannagram** | 0.9999 | 1.0000 | **1.0000** |
| | Cactus | 0.9960 | 0.9999 | 0.9980 |
| | Mauve | 0.7756 | 0.7765 | 0.7761 |
| Inversions, high | **pannagram** | 0.9998 | 1.0000 | **0.9999** |
| | Cactus | 0.9953 | 0.9998 | 0.9976 |
| | **Mauve** | 0.5779 | 0.5787 | **0.5783** |

Table S6: Alignment accuracy on a 32 Mb *Drosophila* chromosome (Chr5; mean of four descendants).

| Variant / divergence | Tool | Recall | Precision | $F_1$ |
|---|---|---|---|---|
| SNP, low | pannagram | 0.9958 | 0.9991 | 0.9974 |
| | Cactus | 0.9999 | 0.9999 | 0.9999 |
| | Mauve | 0.8624 | 0.9996 | 0.9259 |
| SNP, high | pannagram | 0.9966 | 0.9994 | 0.9980 |
| | Cactus | 0.9998 | 0.9998 | 0.9998 |
| | Mauve | 0.8789 | 0.9994 | 0.9353 |
| Inversions, low | **pannagram** | 0.9968 | 0.9994 | **0.9981** |
| | Cactus | 0.9952 | 0.9988 | 0.9970 |
| | Mauve | 0.7989 | 0.9246 | 0.8572 |
| Inversions, high | **pannagram** | 0.9966 | 0.9993 | **0.9980** |
| | Cactus | 0.9901 | 0.9973 | 0.9937 |
| | Mauve | 0.7215 | 0.8387 | 0.7757 |

#### S2.5 Results: *Anopheles* (28.5 Mb chromosome)

The experiment was repeated on *A. coluzzii* Chr1 (28.5 Mb) (Table S7).

Table S7: Alignment accuracy on a 28.5 Mb *Anopheles* chromosome (Chr1; mean of four descendants).

| Variant / divergence | Tool | Recall | Precision | $F_1$ |
|---|---|---|---|---|
| SNP, low | pannagram | 0.9950 | 0.9987 | 0.9968 |
| | Cactus | 0.9968 | 0.9971 | 0.9970 |
| | Mauve | 0.7075 | 0.9985 | 0.8281 |
| SNP, high | **pannagram** | 0.9964 | 0.9994 | **0.9980** |
| | Cactus | 0.9961 | 0.9965 | 0.9963 |
| | Mauve | 0.7218 | 0.9986 | 0.8377 |
| Inversions, low | **pannagram** | 0.9975 | 0.9999 | **0.9987** |
| | Cactus | 0.9821 | 0.9896 | 0.9859 |
| | Mauve | 0.6043 | 0.9014 | 0.7235 |
| Inversions, high | **pannagram** | 0.9965 | 0.9997 | **0.9981** |
| | Cactus | 0.9563 | 0.9758 | 0.9660 |
| | Mauve | 0.5620 | 0.8432 | 0.6744 |

On *Anopheles*, pannagram is the best or tied-best method at every point, including SNPs; Cactus degrades on inversions at high divergence ($F_1 = 0.966$), and Mauve is far behind throughout ($F_1 = 0.67$–$0.84$), losing ≈30% of homologous bases even on SNP-only data.

#### S2.6 Interpretation

Three conclusions hold consistently across all three organisms (*E. coli* 4.6 Mb; *Drosophila* 32 Mb; *Anopheles* 28.5 Mb). (i) On simple substitutions pannagram and Cactus are equivalent ($F_1 \geq 0.997$ everywhere; Cactus marginally ahead on *Drosophila*, pannagram marginally ahead on *Anopheles* at high divergence), whereas Mauve degrades sharply with genome complexity—from $F_1 \approx 0.999$ on *E. coli* to 0.93 (*Drosophila*) and 0.83 (*Anopheles*), losing up to 30% of homologous bases on SNP-only data. (ii) The methods separate most on **structural variation**: pannagram recovers inverted regions near-perfectly and, uniquely, does *not* degrade with divergence (E. coli 1.0000 → 0.9999; Drosophila 0.9981 → 0.9980; Anopheles 0.9987 → 0.9981), and is the best or tied-best on inversions at every point tested; Cactus is strong but slips at high divergence on *Anopheles* ($F_1 = 0.966$); Mauve loses a large fraction of inverted bases everywhere and worsens with divergence (0.58–0.78). (iii) SibeliaZ cannot produce a base-level alignment at these scales at all. Taken with the cost benchmark, pannagram matches the most accurate method (Cactus) on SNPs, is the most accurate on structural variants, aligns a comparable fraction of each genome, and delivers this at 3–13× less runtime and roughly one third to one tenth of the peak memory.

#### S2.7 Limitations

Truth covers SNPs, short indels and inversions; CNV duplications and translocations are outside the pairwise base-level metric used here. Four descendants per point (no cross-replicate variance on the simulation seed beyond the four descendants). Divergence spans ≈0.5–2%; higher divergence and additional taxa would further stress the methods.

### S3 Computational benchmark against pairwise whole-genome aligners

#### S3.1 Aim and scope

Supplementary Note S1 compares pannagram with reference-free *multiple*-genome aligners. This note addresses the complementary class: *pairwise* whole-genome aligners, against which pannagram is run in its reference-based mode (`-ref`), aligning one query genome to one reference. Seven comparators spanning the main algorithmic strategies were used: seed–chain–extend minimizer aligners (minimap2, unimap), suffix/anchor-based aligners (nucmer/MUMmer4, LAST, GSAlign), a mapping-plus-WFA aligner (wfmash), and an adaptive-seed index aligner (FastGA). Because each tool emits a different product, all are scored on a single common quantity—base-level query coverage—defined below. Cost is reported at both 1 and 16 cores, since single-core cost isolates algorithmic work from parallel efficiency.

AnchorWave was excluded because it requires a reference GFF annotation, which was unavailable for both assemblies; LASTZ was excluded because it is single-threaded by design and does not complete on genomes of this size within a practical budget.

#### S3.2 Datasets

Two intraspecific chromosome-scale pairs were used (Table S8), drawn from the same panels as Supplementary Note S1. The two pairs differ in size, repeat content and divergence: *A. coluzzii* is larger and more divergent/repetitive than *D. melanogaster* (pairwise Mash distances ≈0.015 vs ≈0.005). $L_q$, the total length of the query genome, is the denominator of every coverage value reported here.

Table S8: Benchmark pairs. Both are intraspecific (same species, different assemblies). $L_q$ is the total length of the query genome.

| Species | Reference | Query | Chr. | $L_q$ (bp) |
|---|---|---|---|---|
| *A. coluzzii* | GCA_016097095.1 | GCA_016097175.1 | 5 | 246,793,427 |
| *D. melanogaster* | GCA_000001215.4 | GCA_003397115.2 | 6 | 134,496,082 |

#### S3.3 Hardware and scheduling

All runs were executed on dual-socket Intel Xeon Platinum 8368 @ 2.40 GHz nodes (76 physical cores, 1 TB RAM). Each (tool, species, thread-count) combination was submitted as one independent SLURM job with a cgroup-bound CPU allocation (`--cpus-per-task` equal to the thread count, minimum 2; `--mem=96G`), so that no two measured runs competed for the same cores. Wall-clock time, peak resident set size and CPU utilisation were recorded with `/usr/bin/time -v` wrapping the complete invocation, *including* any index construction performed by the tool itself.

This isolation is not a formality. Identical pannagram runs launched on a shared interactive node rather than as cgroup-bound batch jobs were inflated by 36–76% (e.g. an untouched BLAST stage rose from 303 s to 425 s), enough to invert tool rankings; all timings below are from dedicated allocations. Reproducibility was checked by repeating a full pannagram configuration, which agreed to within ≈1% wall-clock.

##### S3.4 Software versions and commands

Tools were installed in isolated `micromamba` environments and run at default parameters unless stated (Table S9). `T` denotes the thread count (1 or 16).

Table S9: Aligner versions and invocations (abridged).

| Tool | Version | Command |
|---|---|---|
| pannagram (`-ref`) | blastn 2.17.0+, R 4.5.3 | `pannagram -ref ref -nchr N -cores T -part_len 1000 -word_size 28 -word_size_gap 15` |
| FastGA | 1.5 | `FastGA -TT -pafx ref.fa qry.fa` |
| minimap2 | 2.31 | `minimap2 -cx asm10 -t T ref qry` |
| unimap | 0.1 | `unimap -c -x asm10 -t T ref qry` |
| nucmer (MUMmer4) | 4.0.0rc1 | `nucmer -t T ref qry && delta-filter -1` |
| wfmash | 0.7.0 | `wfmash -t T ref.fa qry.fa` |
| LAST | 874 | `lastdb -uNEAR db ref && lastal db qry \| last-split` |
| GSAlign | 1.0.22 | `GSAlign -r ref -q qry -t T -fmt 1` |

##### S3.5 Coverage definition

To compare heterogeneous outputs (PAF, MAF, delta, HDF5) on one axis we use *base-level query coverage*: the number of query-genome positions assigned a reference position by at least one alignment, divided by $L_q$,

$$\text{coverage} \;=\; \frac{\left|\bigcup_{a\in\mathcal{A}} Q(a)\right|}{L_q},$$

where $Q(a)$ is the set of query positions inside `M/=/X` CIGAR operations of alignment $a$; insertions consume query but carry no reference position and are excluded. Overlapping and multi-mapping alignments are counted once (set union), so the value cannot exceed 100%.

Three format-specific points matter for reproducibility. (i) For nucmer the one-to-one filtered delta (`delta-filter -1`) must be used; the unfiltered delta double-counts redundant alignments and yields values above 150%. (ii) For pannagram, coverage is the count of non-zero entries of the final HDF5 position matrix (`/accs/qry`), which is by construction a one-to-one query-to- reference map. (iii) FastGA emits its *first* input genome in the PAF query column, the opposite convention to the other tools; its coverage was therefore computed on the target (6th) column. Two independent implementations of the coverage computation agreed exactly on every output.

##### S3.6 Results

Table S10 reports wall-clock time at 1 and 16 cores, peak memory at both, and query coverage.
**Notes.** (i) GSAlign terminated abnormally when restricted to a single core on both species (exit code 1 and segmentation fault, respectively) while completing normally on 16 cores; single-core entries are reported as "—". (ii) The bioconda build of LAST is compiled without multi-threading, so its 1- and 16-core times are equal within noise and it is the slowest tool at 16 cores. (iii) All tools ran at default parameters; FastGA's default identity threshold is permissive ($-i$ 0.7, i.e. 70%), which contributes to its high coverage. (iv) pannagram's single-core peak memory (4.95 GB on *Anopheles*) exceeds its 16-core value (0.60 GB); this arises from a large transient object on the serial gap-merging path and does not affect the 16-core figures.

Table S10: Pairwise whole-genome alignment: wall-clock time, peak memory and base-level query coverage at 1 and 16 cores. Rows ordered by 16-core time. Best value per column and species in **bold**.

| | | Time (s) | | Peak RAM (GB) | | |
|---|---|---|---|---|---|---|
| Species | Tool | 1 core | 16 cores | 1 core | 16 cores | Coverage (%) |
| *A. coluzzii* | FastGA | **167** | **36** | 0.92 | 1.00 | **95.44** |
| | unimap | 255 | 74 | 6.96 | 10.58 | 78.08 |
| | minimap2 | 328 | 81 | 2.49 | 7.71 | 81.96 |
| | wfmash | 1,346 | 103 | 1.19 | 1.98 | 71.83 |
| | pannagram | 777 | 158 | 4.95 | **0.60** | 80.40 |
| | GSAlign | — | 183 | — | 2.15 | 83.29 |
| | nucmer | 1,647 | 626 | 2.94 | 3.87 | 85.79 |
| | LAST | 1,336 | 1,439 | 3.58 | 3.58 | 93.74 |
| *D. melanogaster* | FastGA | **76** | **18** | 0.61 | 1.14 | **97.55** |
| | unimap | 118 | 32 | 6.14 | 8.60 | 93.03 |
| | minimap2 | 147 | 36 | 1.62 | 5.07 | 93.42 |
| | wfmash | 552 | 68 | 0.69 | 1.58 | 77.11 |
| | pannagram | 303 | 77 | 2.51 | **0.26** | 92.87 |
| | GSAlign | — | 82 | — | 1.38 | 92.69 |
| | nucmer | 571 | 155 | 1.65 | 2.35 | 95.03 |
| | LAST | 668 | 688 | 1.18 | 1.18 | 96.76 |

#### S3.7 Observations

Peak memory is again the clearest differentiator, and it separates pannagram from every comparator: at 16 cores pannagram used 0.26 GB (*Drosophila*) and 0.60 GB (*Anopheles*), **4–40× less** than the other aligners (1.2–10.6 GB), while its coverage (92.9% / 80.4%) exceeds wfmash and is comparable to minimap2, unimap and GSAlign. This follows directly from the chunked-BLAST design: pannagram never holds a whole-genome index in memory, whereas the minimizer aligners must (unimap peaks at 10.6 GB). The practical consequence is that pannagram runs on commodity hardware where index-resident aligners do not.

On speed, three regimes appear. FastGA is in a distinct class, 2× faster than the next tool at 16 cores *and* attaining the highest coverage on both species; its adaptive-seed index avoids the seed–chain–extend cost model entirely. LAST and nucmer buy high coverage (93.7–96.8% and 85.8–95.0%) at a large time cost, LAST additionally handicapped by the absence of threading. pannagram is mid-range on both axes. Notably the coverage ranking is not simply a function of runtime: wfmash is fast at 16 cores yet aligns the least (71.8–77.1%), while pannagram is slower than minimap2 but aligns a comparable fraction.

Comparing the two species also separates algorithmic cost from parallel efficiency. Every tool is markedly cheaper on *Drosophila* (smaller, less repetitive, less divergent), but the gap between 1 and 16 cores varies widely: pannagram scales 4.9×/3.9× and FastGA 4.6×/4.2× over 16 cores, whereas LAST does not scale at all.

#### S3.8 Parameter sweep: the two BLAST word sizes

pannagram's `-ref` pipeline invokes `blastn` at two independent stages: step 3 aligns fixed-length query *parts* against the reference, and step 7 re-searches the *gaps* between the resulting synteny blocks. These are controlled by separate word sizes, `-word_size` ($w$, step 3) and `-word_size_gap`

($w_g$, step 7). A full factorial sweep $w \in \{15, 20, 28\} \times w_g \in \{11, 15, 20, 28\}$ was run on both species at one core (Tables S11, S12); these measurements are the basis of the default values used throughout this work.

Table S11: Single-core parameter sweep, *A. coluzzii*. Per-step wall-clock seconds for the dominant stages: step 3 (parts BLAST), step 7 (gaps BLAST), step 8 (gap merging), step 9 (reference matrix assembly). Ordered by total time.

| $w$ | $w_g$ | Total (s) | Step 3 | Step 7 | Step 8 | Step 9 | Coverage (%) | |
|---|---|---|---|---|---|---|---|---|
| 28 | 28 | 729 | 185 | 254 | 127 | 40 | 80.060 | |
| 28 | 20 | 750 | 187 | 260 | 141 | 40 | 80.277 | |
| 28 | 15 | 799 | 186 | 288 | 164 | 41 | 80.399 | ← **default** |
| 20 | 28 | 846 | 300 | 258 | 127 | 40 | 80.093 | |
| 20 | 20 | 866 | 298 | 265 | 140 | 40 | 80.307 | |
| 20 | 15 | 923 | 300 | 292 | 170 | 40 | 80.429 | |
| 28 | 11 | 1,011 | 182 | 437 | 233 | 39 | 80.619 | max. coverage |
| 20 | 11 | 1,135 | 299 | 445 | 231 | 39 | 80.643 | previous default |
| 15 | 28 | 1,275 | 740 | 255 | 120 | 39 | 80.097 | |
| 15 | 20 | 1,300 | 740 | 260 | 139 | 41 | 80.310 | |
| 15 | 15 | 1,346 | 735 | 286 | 164 | 40 | 80.432 | |
| 15 | 11 | 1,560 | 739 | 428 | 230 | 40 | 80.645 | |

Table S12: Single-core parameter sweep, *D. melanogaster*. Columns as in Table S11.

| $w$ | $w_g$ | Total (s) | Step 3 | Step 7 | Step 8 | Step 9 | Coverage (%) | |
|---|---|---|---|---|---|---|---|---|
| 28 | 28 | 278 | 66 | 90 | 32 | 22 | 92.790 | |
| 28 | 20 | 289 | 69 | 90 | 37 | 24 | 92.846 | |
| 28 | 15 | 291 | 68 | 92 | 39 | 23 | 92.873 | ← **default** |
| 20 | 15 | 305 | 87 | 91 | 37 | 22 | 92.875 | |
| 20 | 28 | 305 | 91 | 86 | 35 | 23 | 92.793 | |
| 20 | 20 | 311 | 92 | 87 | 37 | 24 | 92.848 | |
| 28 | 11 | 354 | 70 | 131 | 61 | 23 | 92.903 | max. coverage |
| 20 | 11 | 369 | 85 | 134 | 60 | 22 | 92.904 | previous default |
| 15 | 28 | 431 | 223 | 88 | 32 | 22 | 92.795 | |
| 15 | 20 | 434 | 223 | 91 | 34 | 22 | 92.850 | |
| 15 | 15 | 442 | 222 | 93 | 37 | 23 | 92.877 | |
| 15 | 11 | 493 | 220 | 129 | 56 | 22 | 92.906 | |

**Coverage is nearly invariant across the grid.** Over the full $3 \times 4$ sweep coverage spans only 80.06–80.65% (*Anopheles*; 0.58 percentage points, pp) and 92.79–92.91% (*Drosophila*; 0.11 pp), while total run time varies more than two-fold (729–1,560 s and 278–493 s). Two mechanistically distinct effects explain this.

**(i) The parts word size $w$ is coverage-neutral and should be maximised.** At fixed $w_g$=11, coverage is identical within noise for $w = 15/20/28$ (80.645 / 80.643 / 80.619% on *Anopheles*; 92.906 / 92.904 / 92.903% on *Drosophila*), yet step 3 falls from 739 s to 299 s to 182 s. Coarser seeding at step 3 does lose part-level hits, but those losses are recovered by the downstream gap-filling stage and therefore never reach the final alignment. Hence $w$=28 is an essentially free $\approx 2\times$ speed-up of step 3, and $w$=15 is strictly dominated: 2.5× slower for no coverage gain.

**(ii) The gap word size $w_g$ is a shallow, asymmetric trade-off.** Lowering $w_g$ from 28 to 11 buys only ≈0.1–0.3 pp of coverage per step on *Anopheles* (≈0.02–0.06 pp on *Drosophila*) but is expensive: at $w$=28, moving from $w_g$=15 to $w_g$=11 costs 288 → 437 s in step 7 *and* 164 → 233 s in step 8. The step 8 penalty is the informative part: a more sensitive gap search does not merely search harder, it emits far more candidate hits (≈2.5 million genome-wide at $w_g$=11), the majority of which are redundant overlaps subsequently discarded by greedy chaining during merging. The cost of $w_g$=11 is thus dominated by flooding a downstream stage rather than by the search itself.

**Chosen defaults.** $(w, w_g) = (28, 15)$ was adopted, at the knee of the speed/coverage curve. Relative to $(20, 11)$ it reduces single-core time by **30%** on *Anopheles* (1,135 → 799 s) and **21%** on *Drosophila* (369 → 291 s), for 0.24 pp and 0.03 pp of coverage respectively. Where maximal coverage is required, $(28, 11)$ retains the coverage of the previous default while still being faster (1,011 vs 1,135 s). Both remain user-tunable (`-word_size`, `-word_size_gap`).

#### S3.9 Implementation optimisation of the reference pipeline

Independently of parameter tuning, three implementation-level changes were made to the `-ref` pipeline; all were verified to leave the output byte-identical.

1. **Step 7 — removal of per-batch database construction.** The gap BLAST built one `makeblastdb` database per gap batch (≈1,000 batches genome-wide), each consumed by exactly one `blastn` call. As each database is tiny and used once, this was pure per-invocation overhead (≈0.1 s per call, measured). Replacing `-db` with `blastn -subject`, which searches a subject FASTA directly, removes ≈1,000 processes per run; hit-level output was verified identical on real gap batches (24,487 vs 24,487 hits over 12 batches, zero differing rows).
2. **Step 8 — removal of a quadratic sequence concatenation.** The merge stage glues exactly-adjacent collinear blocks. The original code concatenated aligned sequences pairwise inside the merge loop, re-copying a growing accumulator at each merge, so gluing a chain of $K$ blocks copied $O(K^2)$ characters. Because glue decisions depend only on block coordinates and never on the sequences, the loop was split into a coordinate pass recording the merge topology and a single concatenation per resulting chain, giving $O(n + \sum \text{length})$. Output is byte-identical (300 randomised cases plus real data); a 4,000-block chain improves $> 500\times$.
3. **Step 9 — byte-level gap masking.** Alignment gap positions were obtained by splitting each aligned sequence into a character vector. Replacing this with a raw-byte comparison (byte-identical for ASCII sequence) is $\approx 11\times$ faster and reduced step 9 from 54 s to 41 s.

Together these reduce single-core *Anopheles* run time from 1,222 s to 799 s (**1.53**×) and 16-core time from 221 s to 158 s, at a cost of 0.24 pp coverage.

#### S3.10 Limitations

(i) All measurements are single runs ($N$=1) per configuration; reproducibility was established separately (≈1% wall-clock agreement on a repeated pannagram configuration), and the cgroup isolation removes the dominant source of timing noise, but no confidence intervals are reported. (ii) All tools were run at default parameters; each could likely be tuned further, and FastGA's

Table S13: Cumulative effect of the optimisations, single-core *A. coluzzii*. The implementation changes are output-preserving; only the word-size retuning alters coverage.

| Configuration | Total (s) | Coverage (%) |
|---|---:|---:|
| Baseline, $(w, w_g) = (20, 11)$ | 1,222 | 80.64 |
| + `-subject`, $O(n)$ block gluing, byte-level masking | 1,123 | 80.64 |
| + retuned word sizes $(28, 15)$ | **799** | 80.40 |

high coverage partly reflects a permissive default identity threshold (70%). (iii) Both pairs are intraspecific and of low divergence, a regime favouring $k$-mer/minimizer seeding; behaviour at interspecific divergence is not addressed. (iv) Coverage measures how much query sequence is placed, not whether it is placed *correctly*; alignment accuracy against a known ground truth is assessed separately in Supplementary Note S2. (v) GSAlign single-core measurements are missing because the tool crashed in that configuration.

### S4 Pangenome mobile-element families vs de-novo TE annotation (EDTA)

**Design.** pannagram derives mobile-element families (MEFs) from the *polymorphic* structural variants of a reference-free pangenome, whereas EDTA annotates transposable elements (TEs) de-novo in single genomes. We asked what the pangenome view recovers that EDTA does not, on the five-genome *D. melanogaster* panel. pannagram (`features -sv_families`) grouped 771 large SVs into **57 MEFs**; EDTA v2.3 was run on all five genomes (union library: 2,866 TE consensi). A MEF member was counted as present in EDTA if it (i) matched the EDTA library by BLAST ($\geq$50% length, $e < 10^{-5}$) or (ii) overlapped an EDTA `TEanno` feature on the reference by $\geq$50%. Members absent by *both* criteria define EDTA-missed MEFs.

**Result.** By sequence, 348/771 (45%) members had no EDTA library match; by coordinate, 371/771 (48%) had no EDTA annotation overlap; 321/771 (42%) were absent by both. At family level, **19 of 57 MEFs** are absent from the EDTA de-novo annotation by both criteria. Cross-checking these against the canonical *D. melanogaster* TE set (Bergman lab, 127 named families) shows they are overwhelmingly *known* elements (Table S14): **17/19** map to named D. mel TEs, strongly enriched for **non-LTR retrotransposons** (LINE superfamilies Jockey, I, R1, Loa) and DNA transposons (Tc1-Mariner: FB, Bari1, S-element)—the classes structural de-novo annotators resolve worst—plus two LTR/Gypsy elements (HMS-Beagle, Tirant). Only two short families ($\leq$300 bp) remain unclassified.

**Control: the miss is a library gap, not undetectability.** Running RepeatMasker with the canonical library over the same sequences detects **263/270** EDTA-missed members as their named TE at low divergence (e.g. family 39 = *baggins*, LINE/Loa, 3.8–9.1% divergence; invisible to `blastn` but clear to RepeatMasker). Thus these elements are not undetectable by homology—EDTA missed them because its *de-novo* library was incomplete for these (mostly LINE) families. The fair reading is therefore about de-novo discovery: from pangenome polymorphism pannagram recovers real, canonical, polymorphic TEs—predominantly non-LTR LINEs—that EDTA's de-novo pipeline failed to model, without any pre-existing curated library.

Table S14: The 19 MEFs absent from EDTA's de-novo annotation (by both sequence and coordinate), with their best-matching canonical *D. melanogaster* TE. Names from nucleotide (BLAST) and, for divergent copies, protein-level (`tblastx`) / RepeatMasker matches.

| Family | Copies | Canonical TE | Class |
|---:|---:|---|---|
| 2 | 65 | jockey | LINE/Jockey |
| 5 | 43 | Doc | LINE/Jockey |
| 9 | 43 | F-element | LINE/Jockey |
| 8 | 18 | FB | DNA/Tc1-Mariner |
| 13 | 14 | I-element | LINE/I |
| 6 | 12 | BS | LINE/Jockey |
| 19 | 11 | Juan | LINE/Jockey |
| 40 | 10 | S-element | DNA/Tc1-Mariner |
| 46 | 8 | HMS-Beagle | LTR/Gypsy |
| 16 | 6 | Bari1 | DNA/Tc1-Mariner |
| 23 | 6 | Rt1a | LINE/R1 |
| 38 | 6 | Ivk | LINE/I |
| 53 | 6 | Tirant | LTR/Gypsy |
| 32 | 5 | I-element | LINE/I |
| 49 | 4 | G6 | LINE/Jockey |
| 54 | 3 | Juan | LINE/Jockey |
| 39 | 3 | baggins | LINE/Loa |
| 37 | 4 | unclassified | — |
| 56 | 3 | unclassified | — |

#### S5 Benchmark of `simsearch` against RepeatMasker

We benchmarked pannagram's `simsearch -on_genome` against RepeatMasker (RM 4.2.3, RM-BLAST 2.17.1) for accuracy, robustness to divergence and cost. Both tools take the same TE library and genome and report library occurrences as genomic intervals; RM was restricted to the provided library (`-nolow -no_is -norna`). Crucially we do *not* score one tool against the other's output. Accuracy is measured against **simulated ground truth**: 21 genomes (7 divergence levels 0–30% × 3 replicates), built by implanting ∼1500 copies of the real 816-sequence *A. thaliana* TE library—with controlled substitutions/indels, $5'/3'$ truncation, nesting and target-site duplications—into a homology-free order-5 Markov background (any hit there is a true false positive). Both tools were run across a full identity/ coverage sweep and scored at nucleotide level (Sensitivity, Precision, F1) and at copy level (a truth copy *recovered* if ≥ 50% of its bp are predicted), stratified by divergence and copy kind. On real sequence we avoid scoring against curated TE catalogues, because every public catalogue is itself seeded by homology search (RepeatMasker is the de-facto standard), so recall-of-catalogue is structurally biased in RepeatMasker's favour. Instead we run *both* tools with the same 816-sequence *A. thaliana* library on the *Arabidopsis lyrata* genome (207 Mb; the species diverged ∼6–10 Mya, so the copies are genuinely diverged from the query consensi) and compare the tools *directly* — no ground truth, no filter — which is also pannagram's native use case of searching a library across related genomes.

We make the matched comparison *without* filtering RepeatMasker's output (RM's per-copy coverage cannot be recovered reliably from its `.out`): instead we restrict the *truth* to the population simsearch targets — full-length copies (`kind=copy`, no truncation) — and measure both tools' recall of it, using RepeatMasker raw. On this subset **simsearch and RepeatMasker are statistically indistinguishable** up to 15% divergence (recall 0.993 vs 0.999; mean paired $\Delta = -0.005$, bootstrap 95% CI $[-0.012, +0.001]$, $p = 0.15$; Table S15, Fig. S0/S1), and simsearch precision is $\geq 0.9999$ throughout. Over the *whole* truth (including the truncated / nested / diverged tail) RepeatMasker raw recovers more, because that tail — old, fragmentary copies — is not what simsearch targets: at 0% divergence >97% of simsearch's "misses" are found by BLAST but dropped by the coverage threshold, and lowering `-cov` recovers them at no precision cost (Table S18). This is an *operating-point choice*, not an algorithmic limit. A collinearity-aware HSP-chaining option (`-gap`/`-gapabs`) additionally raises simsearch's own recall of fragmented copies (nested-host fragments 0.70 → 0.76) while keeping the chimeric-merge rate near stock, whereas a naive large gap nearly triples it (Fig. S5); it touches only the vectorised post-BLAST assembly (∼1.5 s, unchanged) and scales to large libraries (rice, 1.5M HSPs, ∼3 min). All of this at 5–7× **lower wall-clock and an order of magnitude less CPU-time** (Table S17, Fig. S4), at comparable memory (∼0.2 GB).

The cross-species comparison reproduces the simulation's conclusions on real, diverged sequence (Table S16). **simsearch is a precision-perfect subset of RepeatMasker**: at the default `sim/cov 85` every base pair it calls, and all 377 of its loci, are also called by RepeatMasker (100% containment)—it never reports anything the community standard would not, direct evidence of precision on real sequence *without* a ground truth. The two tools occupy distinct identity regimes: simsearch's copies have median 95.8% identity to the query consensi (it reports the recent, near-full-length copies), whereas RepeatMasker's span the whole evolutionary spectrum down to ∼60% (median 79.1%)—the operating-point choice of the previous paragraph, now on real cross-species data. It is fully tunable and stays a clean subset: relaxing the thresholds grows simsearch's coverage from 1.2% of RepeatMasker's annotated bp (default) to 16%, 31% and 44% (`sim70/cov70`, `60/50`, `50/25`), always 100% contained in RepeatMasker; it never reaches 100% because RepeatMasker descends past the BLAST cross-species seeding limit (<70% identity), the same cliff seen at high divergence in the simulation. And it does so 9× faster (81 vs 757 s), producing a 379-interval

| | **full-length recall** (truth-restricted) | | | simsearch | simsearch (all truth) | |
|---|---|---|---|---|---|---|
| div. % | simsearch 85 | simsearch 70 | RM raw | precision | Sn | F1 |
| 0 | 1.000 | 1.000 | 0.999 | 0.9999 | 0.892 | 0.943 |
| 5 | 1.000 | 1.000 | 0.999 | 0.9999 | 0.878 | 0.935 |
| 10 | 0.999 | 1.000 | 0.998 | 0.9999 | 0.884 | 0.939 |
| 15 | 0.974 | 0.991 | 1.000 | 0.9999 | 0.873 | 0.932 |
| 20 | 0.027 | 0.943 | 0.999 | 0.9997 | 0.003 | 0.006 |
| 25 | 0.001 | 0.860 | 0.998 | — | 0.000 | 0.000 |
| 30 | 0.000 | 0.331 | 0.997 | — | 0.000 | 0.000 |

Table S15: Simulated ground truth (mean over 3 replicates), no RM filter. *Left:* recall of **full-length** truth copies (the matched population) — simsearch (85) equals RepeatMasker (indistinguishable to 15% divergence, $\Delta = -0.005$, $p = 0.15$); at $\geq$ 20% its default falls below the 85% identity threshold but `sim/cov 70` tracks RM to ∼25%. *Right:* over *all* truth (incl. the truncated/nested tail RM raw also captures) simsearch keeps precision $\geq$ 0.9997; its lower F1 there is the coverage-threshold operating point, not a miss (Table S18). By copy kind (div. 0–15 pooled) simsearch vs RM raw: intact 0.82/1.00, nested 0.82/1.00, host-fragment 0.76/1.00.

annotation against RepeatMasker's 83,129.

Is simsearch missing copies that meet its own 85/85 criterion? We adjudicated this by *direct alignment*, not by parsing RepeatMasker's `.out` (unreliable: near-duplicate consensus names, rows reporting one name with another's coordinates). Taking every RepeatMasker locus with identity $\geq$ 85 (from the `%div` column alone) that simsearch did not call, extracting its genomic sequence, and re-aligning it to the library with a sensitive `blastn` (coverage taken from the consensus length in the FASTA), only **17 of** 14,110 **(**0.1%**) genuinely reach** 85/85; the rest are short high-identity *fragments* (median 5% consensus coverage) that the `-cov` threshold correctly excludes. So at 85/85 simsearch is both precision-perfect and *recall-complete* for genuine full-length copies. The residual few sit at the seeding limit of the default `megablast -word_size 20`; we expose `-word_size`, and lowering it to 11 (the `blastn` task) raises overall yield on *lyrata* from 379 to 456 loci at unchanged precision (100% containment), for a ∼7× runtime cost — worthwhile only for divergent, cross-species searches (on near-identical copies word 20 is faster and loses nothing, so it stays the default).

Finally, the accuracy reported above is set by a deliberate *operating-point choice*, not by a limit of the method. In `-on_genome` mode the BLAST step carries no identity filter, so `-sim`, `-cov` and `-gap` are all applied to the assembled copy and cost nothing to change (the post-BLAST assembly is ∼1.5 s either way). Sweeping them (Table S18) shows that `-cov` is the dominant knob: relaxing it from 85 to 25 raises F1 from 0.945 to **0.999** at 0% divergence and from 0.921 to 0.993 at 20%, *with precision unchanged at* $\geq$ 0.9998 *throughout* — essentially perfect agreement with the truth, at no runtime cost. We nevertheless ship `-sim 85 -cov 85` as the default **by design**: simsearch is meant to report near-full-length copies of the query sequences, and a low `-cov` changes that contract, admitting short partial hits and turning the output into a fragment-level annotation (the regime RepeatMasker's raw mode already occupies). The 0.94 F1 of the default is therefore a definition of the task, not a ceiling: users who want the truncated/fragmentary tail can lower `-cov` and recover it at full precision. (`-sim` matters only near the identity cliff: at 20% divergence `-sim 85` collapses to F1 0.006 while any `-sim` $\leq$ 80 is unaffected; `-gap 10` vs `30` differ in the third decimal.)

| simsearch setting | loci | bp called | contained in RM | % of RM bp |
|---|---|---|---|---|
| `sim85 cov85` (default) | 462 | 0.46 Mb | **100%** | 1.2% |
| `sim70 cov70` | 4,707 | 6.4 Mb | **100%** | 15.9% |
| `sim60 cov50` | 9,666 | 12.6 Mb | **100%** | 31.4% |
| `sim50 cov25` | 15,243 | 17.8 Mb | **100%** | 44.2% |
| median identity of calls: simsearch 95.8% vs RepeatMasker 79.1%; wall 81 vs 757 s | | | | |

Table S16: **Cross-species real-data concordance** (*A. thaliana* 816-seq library on *A. lyrata*, 207 Mb; no ground truth, no filter). Every base pair simsearch calls is also called by RepeatMasker at all thresholds (100% containment), so it is precision-perfect against the community standard on real diverged sequence; relaxing the thresholds tunably expands its coverage of RepeatMasker's annotated landscape from 1.2% to 44%, always as a clean subset. It never reaches 100% because RepeatMasker descends below 70% identity, past the BLAST cross-species seeding limit — the same cliff seen at high divergence in the simulation. RepeatMasker reports 83,129 intervals to simsearch's 379 at the default.

| | simsearch | | RepeatMasker | | |
|---|---|---|---|---|---|
| setting | wall (s) | CPU (s) | wall (s) | CPU (s) | speed-up[a] |
| *Genome size* (816-seq library, 8 threads) | | | | | |
| 5 Mb | 6.0 | 9.6 | 15.5 | 206 | 2.6× |
| 10 Mb | 5.5 | 11.1 | 24.5 | 417 | 4.5× |
| 20 Mb | 13.0 | 18.0 | 60.5 | 929 | 4.7× |
| 40 Mb | 18.0 | 28.6 | 120.0 | 1882 | 6.7× |
| *Threads* (20 Mb genome, 816-seq library) | | | | | |
| 1 | 14.5 | 18.1 | 325.5 | 817 | 22× |
| 4 | 12.5 | 18.1 | 96.0 | 872 | 7.7× |
| 16 | 12.0 | 17.8 | 54.5 | 933 | 4.5× |
| *Library size* (20 Mb genome, 8 threads) | | | | | |
| 100 | 5.5 | 10.8 | 25.0 | 253 | 4.5× |
| 400 | 9.5 | 15.1 | 43.5 | 605 | 4.6× |
| 816 | 12.5 | 17.9 | 60.0 | 929 | 4.8× |
| *Peak RAM* (20 Mb, `/usr/bin/time -v`): simsearch **0.20 GB**, RepeatMasker **0.24 GB** | | | | | |

Table S17: Computational cost (SLURM accounting on exclusive nodes, mean of 2 replicates; peak RAM by `/usr/bin/time -v`). simsearch is 5–7× faster in wall-clock, uses an order of magnitude less CPU-time, and is nearly flat in genome size and thread count, whereas RepeatMasker scales roughly linearly and needs $\geq 8$ threads to close the wall-clock gap. Memory is not a differentiator. The chaining fix adds no measurable runtime: its collinearity test is vectorised, so post-BLAST assembly stays at ~1.5 s (28k-HSP input).
[a] wall-clock ratio RepeatMasker / simsearch.

| setting | div. % | F1 | precision | copy-recall |
|---|---|---|---|---|
| `-sim 70 -cov 25` | 0 | **0.999** | 0.9998 | 0.993 |
| | 10 | **0.999** | 0.9999 | 0.982 |
| | 20 | **0.993** | 0.9999 | 0.908 |
| `-sim 70 -cov 50` | 0 | 0.993 | 0.9998 | 0.961 |
| | 10 | 0.992 | 0.9999 | 0.955 |
| | 20 | 0.986 | 1.0000 | 0.880 |
| `-sim 70 -cov 70` | 0 | 0.965 | 0.9999 | 0.873 |
| | 10 | 0.961 | 0.9999 | 0.863 |
| | 20 | 0.956 | 1.0000 | 0.802 |
| `-sim 85 -cov 85` | 0 | 0.945 | 0.9999 | 0.828 |
| **(default)** | 10 | 0.933 | 0.9999 | 0.800 |
| | 20 | 0.006[a] | — | 0.015 |

Table S18: **Operating-point sweep** (`-gap 10`, one replicate per divergence, vs simulated ground truth). `-cov` is the dominant knob and it is *free*: the `-on_genome` BLAST carries no identity filter, so all thresholds are applied to the assembled copy and cost no runtime. Relaxing `-cov` to 25 reaches F1 ≈0.999 *with precision unchanged* at ≥ 0.9998. The shipped default `-sim 85 -cov 85` is a deliberate choice to report **near-full-length** copies of the query sequences; a low `-cov` changes that contract, admitting short partial hits and turning the output into a fragment-level annotation. The default's F1 of 0.94 is therefore a definition of the task, not a ceiling.

[a] `-sim 85` collapses once divergence exceeds the identity threshold; any `-sim` ≤ 80 is unaffected (F1 0.99 at 20%), so a lower `-sim` is advisable for divergent libraries.
